# Vascular Injury Changes Topology of Vessel Network to Adapt to Partition of Blood Flow for New Arteriovenous Specification

**DOI:** 10.1101/2020.06.09.141408

**Authors:** Kyung In Baek, Shyr-Shea Chang, Chih-Chiang Chang, Mehrdad Roustei, Yichen Ding, Yixuan Wang, Justin Chen, Ryan O’donnelle, Hong Chen, Julianne W. Ashby, Julia J. Mack, Xiaolei Xu, Susana Cavallero, Marcus Roper, Tzung K. Hsiai

**Author notes:** Corresponding Author: Tzung K. Hsiai, M.D., Ph.D., Departments of Medicine and Bioengineering, David Geffen School of Medicine, University of California, Los Angeles, 10833 Le Conte Ave., CHS17-054A, Los Angeles, CA 90095-1679.

## Abstract

Within vascular networks, wall shear stress (WSS) modulates endothelial cell proliferation and arteriovenous specification. Mechano-responsive signaling pathways enable vessels within a connected network to structurally adapt to properly partition blood flow between different parts of organ systems. Here, we study vascular regeneration in a zebrafish model system, performing tail amputation of the Dorsal Aorta (DA)-Posterior Cardinal Vein (PCV) embryonic circulatory loop (ECL) at 3 days post fertilization (dpf). Following severing the ECL, the topology of the micro-circular network is reorganized to engender local increase in blood flow and peak WSS in the closest Segmental Artery (SeA) to the amputation site. Remodeling of this artery increases its radius, and blood flow. These hemodynamic WSS cues activate post-angiogenic Notch-ephrinb2 signaling to guide network reconnection and restore microcirculation. Gain- and loss-of-function analyses of Notch and ephrinb2 pathways, manipulations of WSS by modulating myocardial contractility and blood viscosity directly implicate that hemodynamically activated post-angiogenic Notch-ephrinb2 signaling guides network reconnection and restore microcirculation. Taken together, amputation of the DA-PCV loop induces changes in microvascular topology to partition blood flow and increase WSS-mediated Notch-ephrinb2 pathway, driving the new DLAV-PCV loop formation for restoring local microcirculation.

## Introduction

Microvascular dysfunction is well-recognized to associate with cardiometabolic disorders, including diabetes, hypertension, hyperlipidemia, and obesity.^1,2^ Healthy microvascular networks achieve complex feats of traffic control: perfusing tissues spread throughout the body, avoiding hydraulic short circuits, and dynamically reallocating fluxes in response to tissue demands. Mechano-responsive signaling pathways enable vessels within a connected network to structurally adapt to properly partition blood flow between different parts of organ systems. Vascular endothelium, the inner lining of arterial or capillary walls, transduces biomechanical wall shear stress (WSS) from blood flow. ^3,4^ In the WSS set-point model, blood vessel growth is responsive to WSS: vessels growth is programmed to produce an increase in radius in vessels with high WSS, or a decrease in radius in vessels with low WSS. Since damaged vessels carry no flow and WSS is nearly absent, this mechanism has no evident role in network repair following vascular injury. Whether and how vascular injury-mediated changes in hemodynamics facilitate vascular network regeneration remains an unexplored question.

Endothelial Notch is a well-known mechano-sensitive signaling pathway.^5^ In response to hemodynamic shear forces, proteolytic activation of Notch receptors (Notch1-4) releases Notch Intracellular Cytoplasmic Domain (NICD) that transmigrates to the nucleus.^6^ NICD cooperates with DNA binding protein Lag-1 (CSL), mastermind-like protein 1 (MAML1) and recombination signal-binding protein for immunoglobulin J region (Rbp-Jκ) for the transcription of the target proteins including Hairy and enhancer of split-1 (*Hes1*) and ephrinb2.^7^ During cardiac morphogenesis, WSS induces the Notch1-ephrinb2-neuregulin 1/erb-b2 receptor tyrosine kinase 2 signaling pathway to initiate cardiac trabeculation.^8,9^ Pulsatile and oscillatory WSS differentially regulate Delta/Serrate/Lag ligands in vascular endothelium and smooth muscle cells.^10^

The transmembrane ephrinb2 ligand mediates developmental angiogenesis and arteriovenous specifications. Targeted disruption in ephrinb2 expression induces arteriovenous malformation with defective sprout formation, and homozygous ephrinb2 knockout arrests primitive arterial and venous vessel growth.^11-13^ Cell-cell contact mediates Notch-ephrinb2 bidirectional signaling to EphB4 receptor tyrosine kinase.^14,15^ Reciprocal expression of ephrinb2/EphB4 allow the dorsal aorta (DA) and posterior cardinal vein (PCV) to undergo arteriovenous specification in zebrafish embryos.^16,17^ Repulsive forward EphB4 signaling promotes sprout angiogenesis and cellular intermingling during capillary boundary formation.^18^

Using the zebrafish (*Danio rerio*) model of tail amputation,^19-21^ we sought to demonstrate local hemodynamic cues and endothelial Notch-ephrinb2 signaling pathway that guide network regeneration following vascular injury. In the microvascular network, the DA carries the blood flow to a parallel network of segmental arteries (SeA) that connect to the dorsal longitudinal anastomotic vessel (DLAV). Segmental veins (SeV) carry the venous blood to the principal cardinal vein (PCV), and the DA connects with PCV to form an embryonic circulatory loop (ECL) in the caudal network. We found that tail amputation, severing the ECL, is followed by localize rerouting of blood to the adjacent SeA, with a local increase in blood flow. Increase in WSS in the vessel is accompanied by an elevated activity of the Notch reporter, *tp1*. Genetic and pharmacologic manipulations to alter hemodynamics implicate that changes in WSS in this vessel with induction of Notch signaling guide arterial specification of DLAV, connecting with PCV to form a new DLAV-PCV loop. In response to tail amputation, our results show that WSS-mediated Notch signaling systematically reconstitute arterial network to restore microcirculation by increasing endothelial ephrinb2 expression and differentiation of new vessels. More generally, our analysis shows that the geometry of microvascular networks allow damage in the network to be followed by localized increases in flow and WSS, activating Notch-ephrinb2 signaling to continue to function beyond angiogenesis, for network reestablishment following injury.

## Results

### Tail amputation increases peak WSS in the SeA closest to the amputation site

To assess the effect of amputation on microvascular flow, we used the double transgenic *Tg(fli1:gfp; gata1:ds-red)* line to simultaneously visualize the vascular network (*gfp*^*+*^) and blood cells (*ds-red*^*+*^). Prior to tail amputation, arterial blood passes through the dorsal aorta (DA) either to a set of parallel SeAs, that drain into the dorsal lateral anastomotic vessel (DLAV), and then into the PCV via SeVs,^22^ or directly to the principal cardinal vein (PCV) via an anastomosis at the caudal end of the trunk^23^ (**Figures 1A, J**). Following tail amputation at 3 days post fertilization (dpf), hemodynamic changes in the distal SeAs and SeVs were evaluated for 4 consecutive days (**Figure 1B**). We used custom-written tracking software (see Methods) to track the individual *ds-red*^*+*^ and assessed the time-average velocity of blood flow and WSS in the amputated region (n=3).^22^ Tail amputation severed circulation through ECL, causing *ds-red*^*+*^ to be routed through nearby SeAs. In particular, viscosity increased 3.6-fold in the SeA closest to the amputation site, but only 2.2-fold in the third closest SeA (**Figures 1D, E, K, Supplementary Movie 1**), whereas velocity remained unchanged throughout the trunk (**Figure 1F**). Average WSS in Se vessels decreases from head to tail, since the integrated WSS for a vessel is equal to the pressure drop across the vessel and its cross-section area. Cross-section areas vary little between vessels, and pressures decrease along the DA. However, *ds-red*^*+*^ and SeA radii closely conform, thus WSS near *ds-red*^*+*^ are large (**Figure 1G**), and our modeling shows that passage of a *ds-red*^*+*^ along a SeA is accompanied by a traveling pulse of high WSS. We account for the heterogeneous WSS by the *peak shear stress portion*, the fraction of time during which the WSS exceeds a threshold. We found that the increase in viscosity produced an increase in *peak shear stress portion* but only within this SeA (**Figures 1H, I**). Increased flow in the proximal SeA triggers two forms of network plasticity: remodeling of the SeA, and formation of new vessels from the DLAV that reconnect DA and PCV. We saw that the SeA radius increased over 1 day post tail amputation (dpa) but did not change further (**Supplemental Figure. 1**). This finding is consistent with the WSS set point model predicting that when the WSS in a vessel is larger than its set point, its radius will increase until the stress set point is reattained. By 4 dpa, a new lumenized vascular loop formed between DLAV and PCV (**Figure 2G**). Thus, we demonstrate that these topological changes are directly linked to the WSS changes within the SeA.

**Figure 1.**
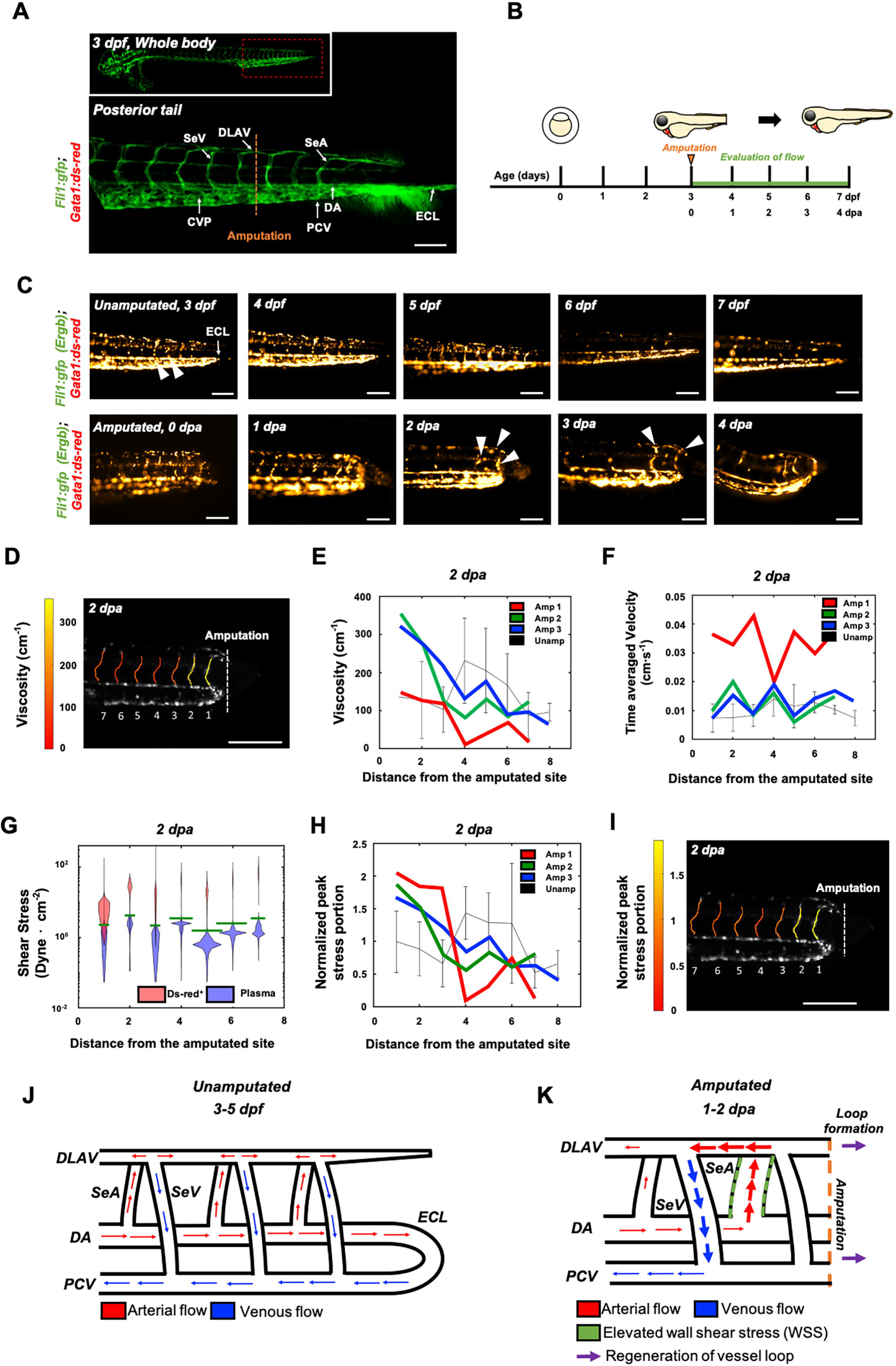
Tail amputation increased peak wall shear stress (WSS) in the segmental artery (SeA) closest to the amputation site. **(A)** Anatomy of tail vasculature in *Tg(fli1:gfp; gata1:ds-red)* embryos; SeA: Arterial segmental vessel, SeV: Venous segmental vessel, DLAV: Dorsal longitudinal anastomotic vessel, PCV: Posterior cardinal vein, DA: Dorsal aorta, CVP: Caudal vein capillary plexus, ECL: Embryonic circulatory loop. Scale bar: 100 μm. **(B)** Experimental design: At 3 days post fertilization (dpf), embryos were randomly chosen for tail amputation (∼100 μm of posterior tail segment). Hemodynamic WSS was evaluated for 4 consecutive days. **(C)** Prior to tail amputation (3 dpf, upper left panel, see also 1J), arterial flow in the DA bifurcates perpendicularly to the SeA. In the caudal vascular network, DA forms an ECL with PCV (white arrow), and venous flow in the DLAV and SeV drain into PCV (white arrowheads). Topological changes in the ECL via tail amputation (0 dpa, bottom left panel, see also 1K) severed circulation via ECL and led to concomitant increase in the number of *ds-red*^*+*^ in the proximal SeA between 1-2 dpa (white arrowheads). **(D-E)** At 2 dpa, viscosity (cm^-1^, number of *ds-red*^*+*^ per unit length of SeA) in the amputated site increased significantly compared to unamputated embryos (Black, mean ± standard deviation across the vessels in 3 unamputated embryos). Viscosity in each SeA was averaged over 3.6 mins. **(F)** Time-averaged velocity (cm·s^-1^) remained unchanged. **(G)** The distribution of WSS exerted by *ds-red*^*+*^ (red) is shifted upward compared to that exerted by plasma (purple), giving intermittent rises of WSS as each *ds-red*^*+*^ passes. Although this separation is present in every SeA, the portion of ECs (red area) experiencing WSS from *ds-red*^*+*^ is higher in the proximal SeAs than in the distal SeAs, leading to a higher peak stress portion. The areas scale with but are not equal to the portion of EC experience stress from *ds-red*^*+*^ and plasma. Green line shows the 0.975 percentile of WSS from plasma, which is used in (**H**) as an activation threshold. For the calculation of WSS, see Methods. (**H-I**) The increase in viscosity resulted WSS to exceed activation threshold in the site of amputation. The threshold is set to 0.975 percentile of plasma shear stress across all intersegmental vessels, and ranges from 1.1 to 5.4 dyne·cm^-2^. **(J-K)** Wiring diagrams illustrate amputation-mediated changes in blood flow and hemodynamic WSS.

**Figure 2.**
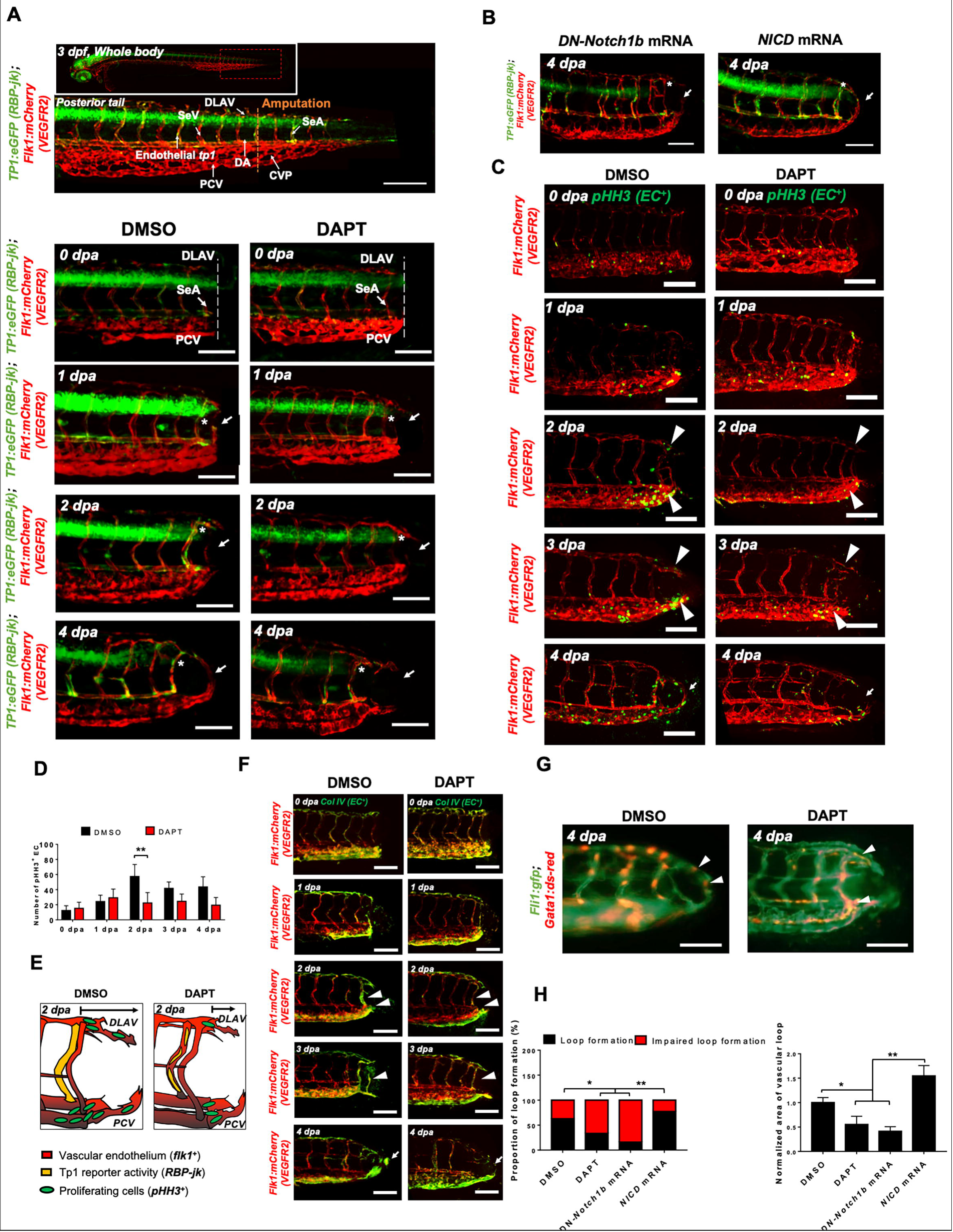
Amputation-mediated WSS induces Notch-dependent DLAV-PCV loop formation. **(A)** Vehicle (DMSO)-treated zebrafish showed prominent endothelial *tp1* activity in the proximal SeA (* asterisk, overlapped yellow) and formed a new vascular loop between the DLAV and PCV at 4 dpa (white arrow). Pharmacological DAPT treatment (100 μM) attenuated endothelial *tp1* activity in the amputated site and neighboring SeA and impaired regeneration of the loop by 4 dpa. (white arrow) (^*\**^ *p < 0.05* vs. DMSO, n=15 per group). SeA: Arterial intersegmental vessel, DLAV: Dorsal longitudinal anastomotic vessel, PCV: Posterior cardinal vein. Scale bar: 20 μm. **(B)** Transient modulation of Notch activity via DN-*Notch1b* and *NICD* mRNAs were performed for positive controls. **(C)** DAPT treatment significantly reduced the number of EC proliferation (*pHH3*^*+*^EC) as compared to DMSO-treated controls at 2 dpa. (white arrowheads, ^*\**^ *p < 0.05* vs. control, n=15 per group) Scale bar: 20 μm. **(D)** Total numbers of endothelial *pHH3*^*+*^ EC in posterior tail segment were assessed to quantify Notch-dependent proliferation. (^*\*\**^ *p < 0.005* vs. DMSO, n=5 per each group) **(E)** Schematic representation of Notch-mediated *pHH3*^*+*^ EC during regeneration. **(F)** *ColIV* expression was prominent in the proximal SeA and regenerated vessels in the DMSO-treated controls (overlapped yellow, white arrowheads). DAPT treatment attenuated *ColIV* expression during regeneration (white arrowheads, n=5 per each group). Scale bar: 20 μm. **(G)** At 4 dpa, endoluminal blood flow was observed in the DMSO-treated embryos (white arrowheads), whereas aberrant blood flow occurred in the amputated site in response to DAPT treatment (white arrowheads). Scale bar: 20 μm. **(H)** Quantification of the proportion of embryos exhibiting loop formation and normalized area of vascular loop (^*\**^ *p < 0.05* vs. DMSO, ^*\*\**^ *p < 0.005* vs. *NICD* mRNA, n= 17 for DMSO, n=20 for DAPT, DN-*Notch1b* and *NICD* mRNAs).

### Tail amputation induced Notch-mediated new DLAV-PCV loop formation

Following tail amputation, transgenic *Tg(tp1:gfp; flk1: mcherry)* embryos displayed prominent endothelial *tp1* activity in the SeA close to the injury site, accompanying the formation of the new vascular loop. Inhibiting Notch signaling by treating with γ-secretase inhibitor (DAPT) attenuated endothelial *tp1* activity in the SeAs and impaired the formation of the DLAV-PCV loop at 4 dpa. (^*\**^ *p < 0.05* vs. DMSO, n=15 per group) (**Figures 2A, F, G**). This local endothelial *tp1* activity and the loop formation was confirmed independently by micro-injecting dominant negative (DN)-*Notch1b* mRNA. Conversely, ectopic overexpression of *NICD* mRNA up-regulated *tp1* activity in the injured SeAs and promoted DLAV-PCV loop formation. (**Figure 2B**). Colocalization of vascular endothelium (*flk1*^*+*^) and cell mitosis marker, phospho-histone 3 (*pHH3*^*+*^), was assessed to evaluate EC proliferation. (**Figure 2C**). While DMSO-treated embryos showed a significant increase in the number of *pHH3*^*+*^ EC in the caudal vein capillary plexus (CVP), DLAV and regenerated vessels, DAPT treatment for 2 days resulted in ∼48% reduction in *pHH3*^*+*^ EC (^*\*\**^*p < 0.005* vs. DMSO, n=5 per group) (**Figures 2C-D**). The number of total *pHH3*^*+*^ cells in the amputated site was also dependent on Notch activation (**Supplemental Figure 3A**). In addition, immunostaining against collagen 4 (*ColIV*) to assess basal lamina deposition showed elevated expression both in the proximal SeA and in regenerated vessels, and this expression was attenuated by DAPT treatment (**Figure 2F**). *In vitro* analyses with human aortic endothelial cells (HAEC) further supported Notch-mediated Matrigel tube formation and endothelial migration for wound recovery. DAPT treatment led to ∼84% reduction in tube length and reduced branch point formation (^*\*\**^ *p < 0.005* vs. DMSO, n=3) (**Supplemental Figures 5A, D, E**). Furthermore, DAPT treatment mitigated HAEC migration and reduced the area of recovery (A.O.R) at 12 hours post scratch (^*\**^ *p < 0.05*, ^*\*\**^ *p < 0.005* vs. DMSO, n=3) (**Supplemental Figures 5B, F**). As a corollary, silencing Notch1 expression with siRNA (*siNotch1*) reduced tube length and branch point formation by ∼40% and ∼50% respectively, and inhibited migration (^*\*\**^ *p < 0.005* vs. *siScr*, n=3) (**Supplemental Figure 5C**). Taken together, our *in vivo* and *in vitro* findings support the role of Notch signaling in EC proliferation and migration for vascular loop formation after injury.

### Changes in WSS modulate DLAV-PCV loop formation in a Notch-dependent manner

To test whether formation of a new vascular loop occurs in response to WSS triggers, we manipulated WSS by transiently modulating zebrafish blood viscosity (since *peak stresses* reflect the flux of *ds-red*^*+*^ through a vessel) and myocardial contractility for microvascular flow (**Figure 3A, Supplemental Figure 2**). Reduction in blood viscosity via *Gata1a* morpholino oligonucleotide (MO) injection or inhibition of myocardial contractility with 2,3-butanedione monoxime (BDM)^33^ impaired DLAV-PCV loop formation at 4 dpa. On the contrary, augmented blood viscosity via erythropoietin (*epo)* mRNA injection or increased contractility via isoproterenol treatment promoted loop formation at 4 dpa. Consistently, increasing plasma viscosity by introducing 6% hydroxyethyl hetastarch via common cardinal vein (CCV) (∼5.5 % volume expansion) also promoted DLAV-PCV loop formation without influencing average heart rate, flow rate of *ds-red*^*+*^ and endogenous expression of Notch-related genes (^*\**^ *p < 0.05*, ^*\*\**^ *p < 0.005*, ^*\*\*\**^ *p < 0.0005* vs. *control* MO, n=20 per each group) (**Figures 3B, Supplemental Figure 6**).^24^ We monitored the effect of the perturbing hemodynamics upon *tp1* activity. At 2 dpa, both *Gata1a* MO injection and BDM treatment reduced endothelial *tp1* activity and the number of *pHH3*^*+*^ EC by ∼42% and ∼65%, respectively, as compared to the *control* MO-injected embryos. Transient *epo* mRNA overexpression or isoproterenol treatment up-regulated endothelial *tp1* activity, and subsequent *pHH3*^*+*^ EC by ∼52% and ∼39%, respectively, at 2 dpa. Injection of 6% hydroxyethyl hetastarch increased endothelial *tp1* activity but did not increase the number of *pHH3*^*+*^ EC at 2 dpa (^*\**^ *p < 0.05*, ^*\*\**^ *p < 0.005*, ^*\*\*\**^ *p < 0.0005* vs. *control* MO, n=5 per each group) (**Figures 3C-E, F-G**). Modulation of both WSS and global Notch activity (DAPT treatment, *NICD* & DN-*Notch1b* mRNA injections) corroborated that WSS-mediated Notch signaling pathway is implicated in DLAV-PCV loop formation (**Supplemental Figure 7**). As a corollary, exposure of HAECs to pulsatile shear stress (PSS, *∂*τ/*∂*t = 71 dyne·cm^2^·s at 1 Hz) up-regulated Notch-related mRNA expression following either DAPT treatment or *siNotch1* transfection (**Supplemental Figure 5G**). Taken together, these results further support that increase in hemodynamic WSS activates endothelial Notch-mediated vascular loop formation.

**Figure 3.**
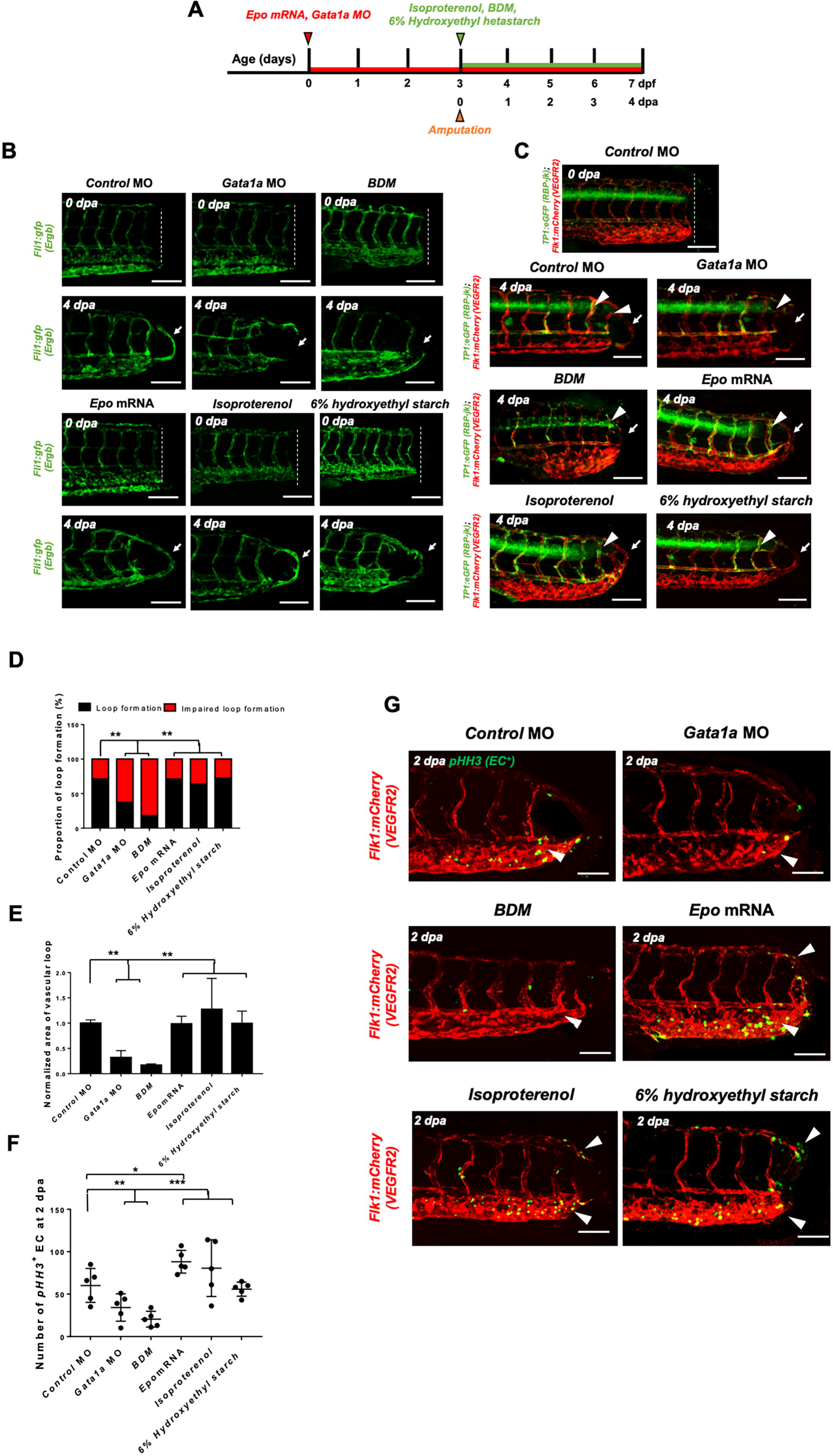
Changes in WSS modulate DLAV-PCV loop formation in a Notch-dependent manner. **(A)** Experimental design to genetically and pharmacologically manipulate hemodynamic WSS in zebrafish embryos. **(B)** *Gata1a* MO injection (1mM) or 2,3-butanedione monoxime (BDM, 100 µM) treatment impaired vascular loop formation at 4 dpa (white arrows). Erythropoietin (*epo*) mRNA injection (10-20 pg/nL), isoproterenol treatment (100 µM) and 6% hydroxyethyl hetastarch promoted regeneration. (^*\*\**^ *p < 0.005* vs. *control* MO, n=20 for each group). Scale bar: 20 μm. **(C)** *Gata1a* MO and BDM treatment reduced endothelial *tp1* activity and impaired regeneration, whereas *epo* mRNA injection or isoproterenol treatment up-regulated endothelial *tp1* activity in the amputated site (white arrowheads) and promoted regeneration at 4 dpa (white arrows) (n=20 per each group). Scale bar: 20 μm **(D-E)** Quantification of the proportion of embryos exhibiting loop formation and normalized area of vascular loop (^*\*\**^ *p < 0.005* vs. *control* MO, n=20 for each group). Scale bar: 20 μm. **(F-G)** Total numbers of endothelial *pHH3*^*+*^ EC in posterior tail were assessed to quantify WSS-dependent EC proliferation. *Gata1a* MO injection or BDM treatment reduced ∼42% and ∼65%, whereas *epo* mRNA and isoproterenol treatment increased *pHH3*^*+*^ EC by ∼52% and ∼39% respectively at 2 dpa. (^*\**^ *p < 0.05*, ^*\*\**^ *p < 0.005*, ^*\*\*\**^ *p < 0.0005* vs. *control* MO, n=5 for each group). Scale bar: 20 μm.

### Partitioned blood flow promotes Notch signaling for arterial specification

To elucidate whether WSS guides arterial specification during the DLAV-PCV loop formation, we utilized the transgenic *Tg (flt1: tdtomatoe; flt4: yfp)* zebrafish line that allows simultaneous visualization of arterial (*flt1*^*+*^) and venous (*flt4*^*+*^) vascular networks (**Figure 4A**). Time-lapse imaging in the presence or absence of DAPT treatment revealed that amputation-mediated Notch signaling regulates *flt1*^*+*^ network during loop formation. DMSO-treated controls developed a secondary loop of *flt1*^*+*^ that extended dorsally from the distal SeA in conjunction with blood flow to anastomose with *flt4*^*+*^ PCV (**Supplemental Videos 2**). Treatment with DAPT inhibited the initiation of *flt1*^*+*^ DLAV to connect with DA (0-1 dpa) and the collateral arterialization of *flt4*^*+*^ DLAV (2-3 dpa),^25^ resulting in an impaired *flt1*^*+*^ network for the loop formation at 4 dpa (**Figure 4B, Supplemental Figure 8**). Reducing viscosity via *Gata1a* MO injection diminished the distal *flt1*^*+*^ in both SeA and DLAV and partially impaired *flt4*^*+*^ DLAV from forming a loop with the PCV **(Figures 4C-D)**. BDM treatment abrogated both *flt1*^*+*^ and *flt4*^*+*^ loop formation. In contrast, both increasing viscosity via *epo* mRNA or 6% hydroxyethyl hetastarch injection or increasing myocardial contractility via isoproterenol treatment promoted *flt1*^*+*^network formation in the amputated site as compared to MO-injected controls **(Figures 4C-D)**. Gain- and loss-of-function analyses of global Notch activity further corroborated that WSS-activated Notch signaling coordinates *flt1*^*+*^*/ flt4*^*+*^ loop formation (n=20 per each group) **(Supplemental Figure 9)**. As a corollary, genetic and pharmacologic elevation of hemodynamic WSS failed to restore loop formation in the presence of DAPT treatment or DN-*Notch1b* mRNA injection; however, *NICD* mRNA injection restored *Gata1a* MO-impaired *flt1*^*+*^/ *flt4*^*+*^ loop formation at 4 dpa (n=20 per each group) **(Supplemental Figure 9).** Taken together, our results indicate that WSS-responsive endothelial Notch signaling reactivates nascent *flt1*^*+*^network and subsequent arterialization of *flt4*^*+*^ DLAV to form a new vascular loop.

**Figure 4.**
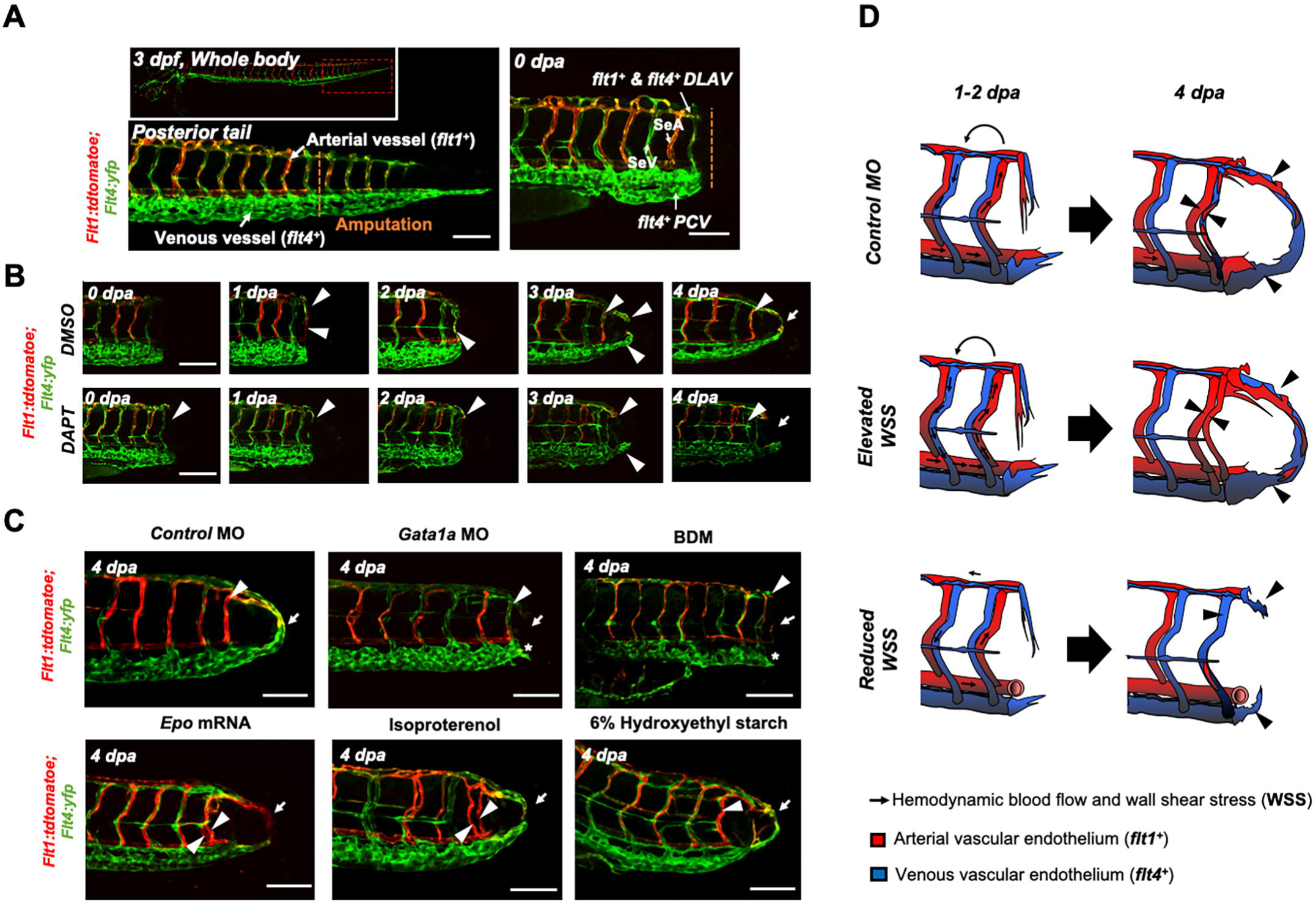
Partitioned blood flow promotes Notch-mediated arterial network. **(A)** A representative image of tail vasculature in the transgenic *Tg(flt1:tdtomatoe; flt4: yfp)* zebrafish line at 3 dpf. *flt1*^*+*^: arterial vascular endothelium, *flt4*^*+*^: venous vascular endothelium. Scale bar: 100 μm. **(B)** In response DMSO treatment, *flt1*^*+*^ preferentially formed an initial dorsal to ventral connection between the DLAV and the DA in the amputated site (white arrowhead). Between 1-2 dpa, *flt4*^*+*^ from the DLAV regenerated toward the DA exhibiting collateral arterial phenotype (overlapped yellow, white arrowheads). While regeneration of *flt4*^*+*^ occurred in the CVP and DLAV for loop formation between 2-3 dpa, the distal segmental network enhanced *flt1* expression at 3 dpa. *flt1*^*+*^ extended dorsally from SeA at 4 dpa and formed a vascular loop with *flt4*^*+*^ regenerated from the PCV (white arrow). Conversely, DAPT treatment inhibited the initial connections of both *flt1*^*+*^ and *flt4*^+^ at 2 dpa and partially inhibited *flt4*^+^ from the DLAV and CVP at 3 dpa (white arrowheads). At 4 dpa, DAPT treatment inhibited *flt1*^*+*^network (white arrowheads) and attenuated loop formation at 4 dpa (white arrows). n=5 per each group. Scale bar: 20 μm. **(C)** Following tail amputation, *Gata1a* MO injection or BDM treatment diminished *flt1*^*+*^ in the amputated site (white arrowheads) and partially attenuated *flt4*^*+*^ from the DLAV and PCV (* asterisk, n=20 per each group). Increase in WSS (*epo* mRNA, isoproterenol, 6% hydroxyethyl hetastarch) enhanced *flt1*^*+*^ network (white arrowheads) during loop formation (white arrows) as compared to MO-injected controls. (n=20 per each group). Scale bar: 20 μm. **(D)** Schematic representations of WSS-mediated arterial- and venous-regeneration. Black arrowheads depict regenerated *flt1*^*+*^ (Red) and *flt4*^+^ network (Blue) in response to differential hemodynamic WSS.

### WSS-responsive Notch-ephrinb2 pathway regulates arterial network formation

To investigate the WSS-responsive downstream signaling pathways underlying DLAV-PCV loop formation, we performed 1) immunostaining for endogenous ephrinb2 and 2) batch processing to assess spatiotemporal variations in endothelial ephrinb2 *in situ* following tail amputation **(Figure 5A)**. Endothelial ephrinb2 staining increased at 1 dpa in the distal DA and DLAV where WSS-responsive arterial network formation occurs. Following the initial anastomosis between the DLAV and DA at 2 dpa, the distal SeA and regenerated vessel region expressed prominent ephrinb2 staining to form a new vascular loop at 4 dpa. *Gata1a* MO injection and BDM treatment attenuated endothelial ephrinb2 staining from 2 to 4 dpa, whereas *epo* mRNA-, isoproterenol-, and 6% hydroxyethyl hetastarch-augmented WSS accentuated staining in the distal SeA, DLAV and CVP from 2 to 4 dpa. DAPT treatment, as the positive control, reduced ephrinb2 staining affirming that the Notch-dependent ephrinb2 pathway is involved in forming the DLAV-PCV loop (n=5 per each group) (**Figure 5B**). Consistent with these observations, PSS induced transient up-regulation of ephrinb2 protein in a time- and Notch-dependent manner in cultured HAEC (n=3 per each time point) (**Supplemental Figure 11A**) As an internal positive control, Kruppel Like Factor 2 (klf2) mRNA expression was also up-regulated (**Supplemental Figure 11B**).^10^

**Figure 5.**
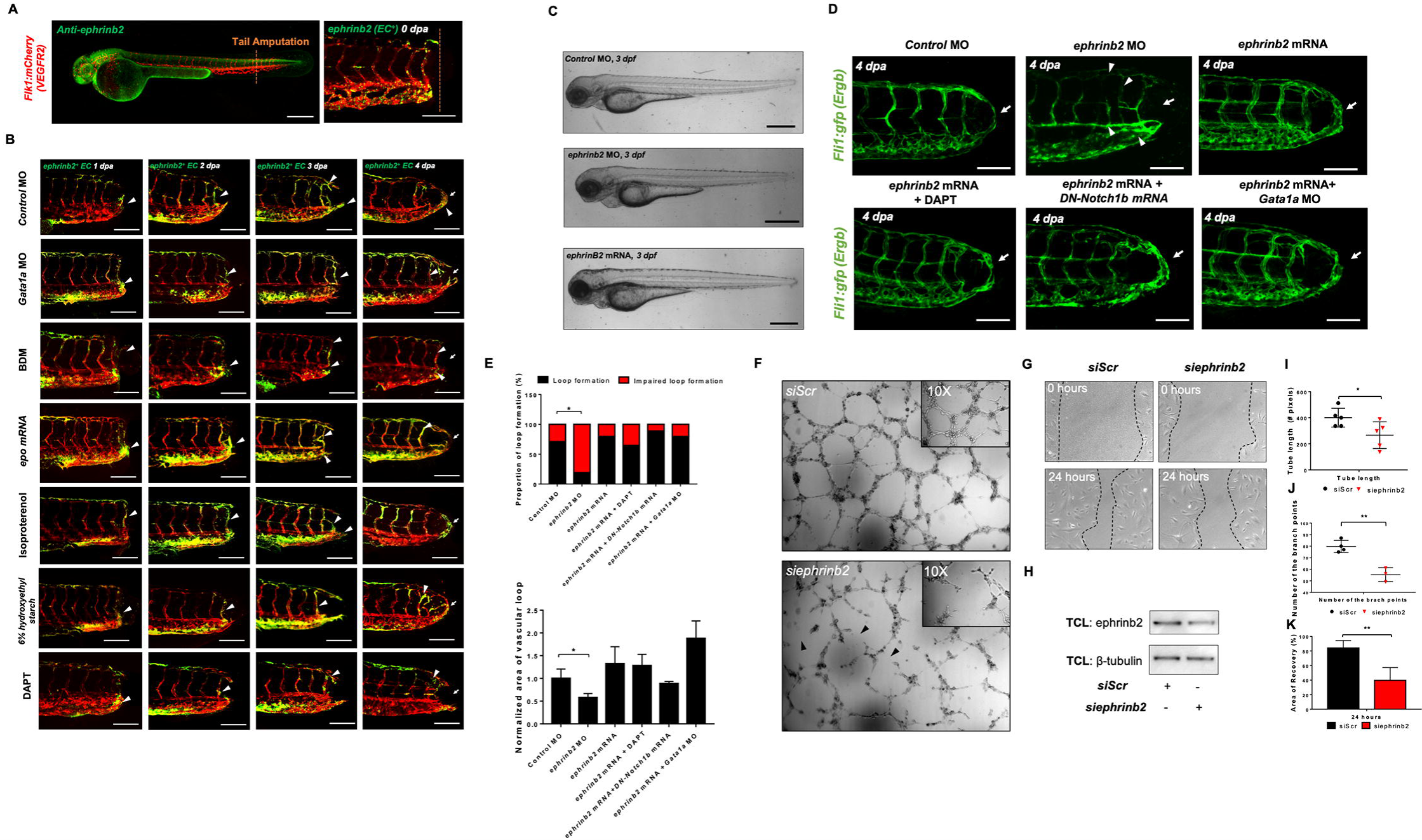
ephrinb2 regulates Notch-dependent vascular loop formation. **(A)** Whole mount immunofluorescence staining against ephrinb2 (green). At 3 dpf, *Tg(flk1: mcherry)* embryos exhibited prominence in endogenous ephrinb2 expression in central nervous system, heart and neutral tube (white arrowheads). Scale bar: 100 μm. **(B)** Representative images of endothelial ephrinb2 staining during DLAV-PCV loop formation (white arrowheads). Scale bar: 20 μm. At 1 dpa, endothelial ephrinb2 staining increased in the caudal DA (white arrowhead). Ephrinb2 staining increased in the distal SeA and regenerated vessels for secondary loop formation at 4 dpa. Reduction of viscosity- and contractility-mediated WSS via *Gata1a* MO injection and BDM treatment reduced endothelial ephrinb2 staining, whereas increase in hemodynamic WSS (*epo* mRNA, isoproterenol, 6% hydroxyethyl hetastarch) accentuated ephrinb2 staining in the distal SeA, DLAV and CVP from 2 to 4 dpa. DAPT treatment, as the positive control, reduced ephrinb2 staining in the distal SeA and regenerated vessel regions (n=5 per each group). **(C**) Gross morphology examinations following global ephrinb2 modulation. Both *ephrinb2* MO and mRNA injections did not affect gross morphology (white arrowheads). Scale bar: 100 μm. **(D)** Compared to the MO-injected controls, ephrinb2 MO injection (0.5-1mM) impaired loop formation (white arrow) followed by retardations in the DA and SV morphology, maturity of the CVP and DLAV at 4 dpa (white arrowheads). Transient overexpression of *ephrinb2* mRNA restored DAPT, DN-*Notch1b* mRNA (10-20 pg/nL) and *Gata1a* MO-impaired loop formation (white arrows). Scale bar: 20 μm. **(E)** Quantification of the proportion of embryos exhibiting loop formation and normalized area of vascular loop (^*\**^ *p < 0.05* vs. *control* MO, n=20 per group). **(F-G)** Representative images of Matrigel and HAEC migration assays following *siScr* or *siephrinb2* transfection. **(H)** The density quantification of ephrinb2 expression following *siephrinb2* transfection. TCL: total cell lysates (**I-J**) *siephrinb2* transfection reduced both tube length and the number of branch points as compared to *siScr*-transfected HAEC. (^*\**^ *p < 0.05*, ^*\*\**^ *p < 0.005* vs. *siScr*, n=3) **(I)** *siephrinb2* transfection reduced A.O.R by ∼45% at 24 hours post scratch (^*\*\**^ *p < 0.005* vs. *siScr*, n=3)

In addition, we genetically manipulated global ephrinb2 expression via MO or mRNA injection to assess loop formation in the transgenic *Tg(fli1: gfp)* zebrafish embryos. Neither knockdown nor overexpression of ephrinb2 expression affected the gross microvascular morphology as compared to the controls (**Figure 5C**). Injection of *ephrinb2* MO impaired the loop formation, followed by collapsed DA morphology and maturity of the DLAV and CVP at 4 dpa. Conversely, transient overexpression of *ephrinb2* mRNA increased endoluminal sizes of regenerated vessels and restored DAPT, DN-*Notch1b* mRNA and *Gata1a* MO-impaired loop formation (^*\**^ *p < 0.05*, ^*\*\**^ *p < 0.005*, ^*\*\*\**^ *p < 0.0005* vs. *control* MO, n=20 per group) (**Figures 5D-E**). As a corollary, *in vitro* Matrigel and migration assays following *siephrinb2* transfection further support these observations. While *siephrinb2* transfection resulted in ∼ 30% and ∼ 32% reduction in the tube length and the number of the branch points, respectively (**Figures 5F, I, J**), A.O.R was reduced by ∼49% at 24 hours post scratch (^*\**^ *p < 0.05*, ^*\*\**^ *p < 0.005* vs. *siScr* transfection, n=3) (**Figures 5 F, K**). In the transgenic *Tg(flt1:tdtomatoe; flt4: yfp)* zebrafish line, injection of *ephrinb2* MO diminished *flt1*^*+*^ in the distal SeA and DLAV and reduced *flt4*^*+*^ in the DLAV and PCV, mimicking phenotypes in the absence of WSS or Notch activity. Transient overexpression of *ephrinb2* mRNA enhanced the dorsal *flt1*^*+*^ network in the presence of DAPT, DN-*Notch1b* mRNA, and *Gata1a* MO and further restored regeneration of *flt4*^*+*^ DLAV and PCV (n=20 per group) (**Figures 6A, B**). Thus, our data corroborate that WSS-responsive Notch-ephrinb2 pathway guide the arterial specification to promote loop formation.

**Figure 6.**
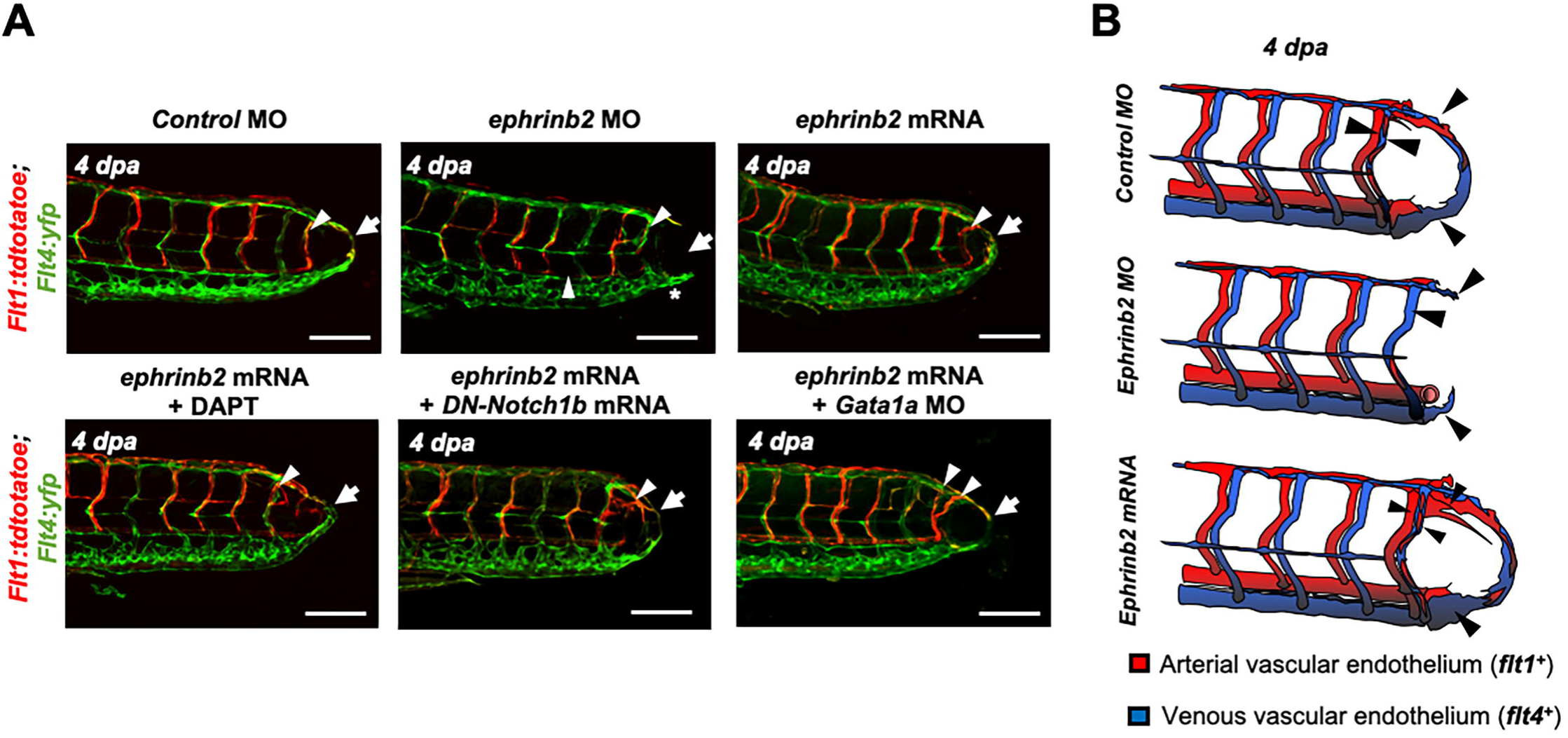
WSS-responsive Notch-ephrinb2 pathway regulates arterial network formation. **(A)** Injection of *ephrinb2* MO (0.5-1mM) diminished *flt1*^*+*^ in the DA and amputated site (white arrowheads) to attenuate loop formation at 4 dpa (white arrow). Transient overexpression of *ephrinb2* mRNA promoted *flt1*^*+*^ network formation in the amputated site (white arrowheads) and restored DAPT and DN-*Notch1b* mRNA-impaired *flt1*^*+*^ regeneration (white arrow) (n=20 per each group). Scale bar: 20 μm. **(B)** Schematic representations of ephrinb2-dependent arterial (*flt1*^*+*^) and venous (*flt14*^*+*^) regeneration. Black arrowheads depict regenerated *flt1*^*+*^ (Red) and *flt4*^+^ (Blue) in response to differential ephrinb2 expressions.

## Discussion

The mechano-sensitive Notch signaling pathway is widely recognized to coordinate vascular proliferation and differentiation.^26-31^ Deletion of Notch1 or Notch4 results in impaired micro-vascular network, whereas EC-specific single allele deletion or pharmacologic inhibition of Delta-like ligand 4 (Dll4) regulates neovascularization.^26,32,33^ Notch signaling is involved in biomechanical bi-potential cell fate decisions,^32^ including, in developing vessels, the trans-differentiation of arterial vessels into veins, and its activity guides the sizes of arteries and veins.^34^ Blood flow-induced endothelial Notch activity systematically remodels the arterial segmental network in developing zebrafish embryos,^35^ and implicates vascular identity during caudal fin regeneration.^6,25^ Our findings demonstrate that peak WSS-activated local Notch activity is also essential to stimulate microvessel regrowth and arterial specification after injury (**Figure 4, Supplemental Figure 9**). Compared to hypoxia, which is commonly implicated in microvascular regrowth,^35,36^ WSS can remain highly localized within vessels near the site of injury to guide arterialization of a new loop for restoring microcirculation.

WSS cues are known to be instrumental in developmental remodeling of vascular networks,^37^ and has also been connected to angiogenic sprouting following injury.^38,39^ However, since injury removes flow carrying vessels, and reduces overall blood supply to the injured tissue, it has not been clear how WSS can increase following injury. Our study shows how by altering the partitioning of blood flow among SeAs, amputation of the ECL loop produces locally elevated WSS. We demonstrate that augmented WSS in the neighboring SeA is a necessary biomechanical cue to up-regulate endothelial Notch-ephrinb2 signaling to drive arterial specification in the DLAV and the growth of a new loop between DLAV and PCV. Mathematical modeling clarified that peak WSS can increase, even when average WSS, which is constrained by the pressure differences in the network, remains constant.

How does tail amputation increase WSS in the proximal SeA? Average WSS in any vessel is constrained by the pressure drop across the vessel, which increases by 57% following amputation of the ECL. However, we found that a much stronger signal is presented by the unsteadiness of the WSS. In the finest vessels, passage of a *ds-red*^*+*^ exerts much larger WSS than plasma, and the peak stress within a vessel is strongly linked to the flux of *ds-red*^*+*^ (**Figure 1**). When micro-vessels bifurcate, *ds-red*^*+*^ and plasma fluxes divide in different ratios: *ds-red*^*+*^ are more likely to enter the larger radius branch of the bifurcation than would be dictated by ratio of flows. Prior to amputation, this effect means that SeAs carry lower viscosity than the DA. But after tail amputation *ds-red*^*+*^ reaching the distal DA must drain through one of the SeAs, causing a large increase in *ds-red*^*+*^ flux, and therefore WSS, in those vessels.

Before amputation, *ds-red*^*+*^ drain directly into the PCV through the ECL. Following amputation, the large increase in peak WSS requires that *ds-red*^*+*^ drain directly into the PCV through SeAs. This ECL emerges very early in embryogenesis,^40^ but direct anastomoses between artery and vein are common in microvascular networks and can form even when fine vessels already connect the two, such as the basal artery in the zebrafish midbrain which drains into the DLAV.^40^ Our results suggest that these loops enable large increases in flow in proximal vessels following amputation, making them directly responsible for initiating WSS-triggered vessel repair and regeneration. Linking vascular regeneration to WSS is particularly relevant to tissues like the embryonic zebrafish trunk, in which oxygen levels are too high for wounding to trigger hypoxic vessel regrowth. ^41^ At the same time, the existence of such loops carries physical costs, since the *ds-red*^*+*^ and glucose carried in the ECL are transported through the trunk without perfusing the trunk tissues.

Endothelial ephrinb2 influences angiogenic sprouting and vessel migration.^42-45^ Ephrinb2 further regulates internalization and subsequent signaling activities of vascular endothelial growth factor receptors (VEGFR2 and VEGFR3), and the expression is prominent at sites of neovascularization or pathological angiogenesis.^46,47^ In response to fluid shear stress, reciprocal expression of ephrinb2/EphB4 determines vascular identity.^48-50^ We observed that endothelial ephrinb2 staining was accentuated in the amputated region, indicating that it plays a role in establishing the new arterial network. Genetic and pharmacological modulations of hemodynamic WSS further supported the notion that augmented WSS is necessary to promote ephrinb2 expression during vascular loop formation. Our data indicate that endogenous ephrinb2 is a primary target underlying the Notch-dependent arterial network in response to tail amputation (**Figure 6**). Ephrinb2 is known to regulate postnatal venous neovascularization by modulating EphB4 expression, and ephrinb2 and EphB4 physically interact in growing vessels; and this interaction is implicated in cellular intermingling and vascular anastomosis.^44,46,51-55^ Following tail amputation, we observed that ephrinb2 and EphB4 expressions exhibited synergistic effects for loop formation (**Supplemental Figure 10A-B**). *In vitro* assays show that expression of both ephrinb2 and EphB4 is necessary for endothelial migration and angiogenic tube formation (**Supplemental Figure 10C-H**). Under hydrodynamic PSS, Notch-dependent ephrinb2 and EphB4 protein expression were up-regulated in a time-dependent manner, and the total level of ephrinb2/EphB4 interaction was increased without influencing the polarization kinetics and distribution of the ephrinb2/EphB4 proximity ligations (**Supplemental Figure 11**). Thus, our observations suggest that amputation-augmented WSS induces arterial Notch-ephrinb2 signaling to regulate lateral venous plexus where EphB4 promotes vessel regeneration. Our proposed mechanism is summarized in **Figure 7**. In contrast to our results which emphasize reconnections from the arterial network, Xu *et al*. report that vein derived arterial cells reconnect vessels following caudal fin regeneration.^56^ The topology of the trunk network causes largest peak WSS increases in Se arteries, and the dominance of arterial regeneration may reflect localization of WSS signals. Whether reconnection begins with the arterial or venous networks may also depend upon the age- or injury-specific conditions.

**Figure 7.**
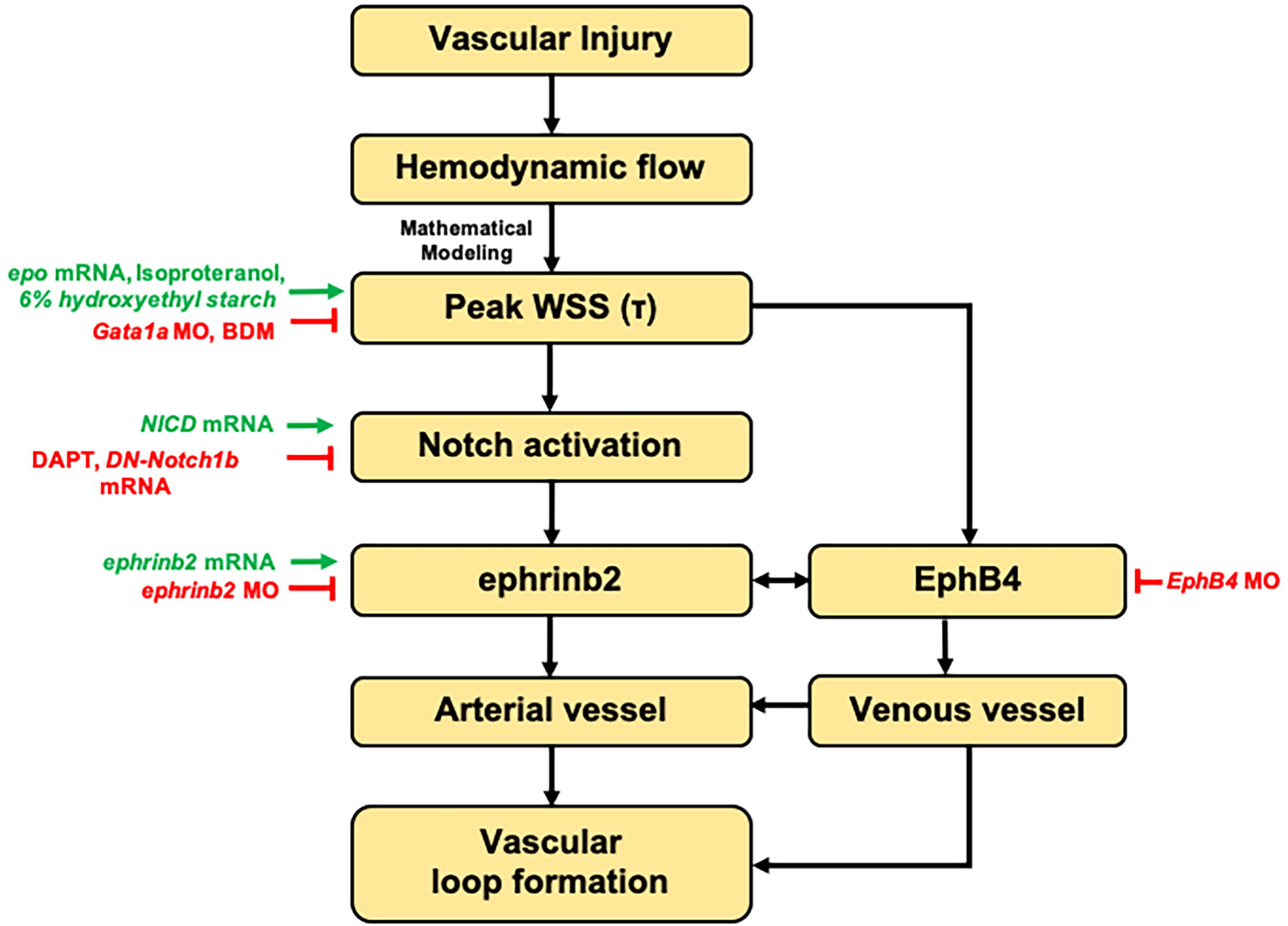
Schematic overviews of the proposed mechanisms.

Our data draw a line from WSS to vascular loop formation through Notch activated ephrinb2/EphB4 expression. However, both WSS and Notch activation are multiply linked to angiogenic cell proliferation and migration, through pathways that may play linked or parallel roles. For example, flow-sensitive microRNAs (miRs), miR-125 and -210, promote tip cell formation and arteriolar branching^57^ in response to ischemic injury or hyperlipidemic stress.^58,59^ Clusters of miR-497∼195 or -449 are implicated in angio- and multicillogenesis via Notch activation.^60,61^ miR-17 and homologs of miR-20 are associated with endothelial ephrinb2 expression.^62^ Both WSS and Notch-activation are connected to angiogenesis-affecting metabolic changes. WSS affects endothelial VEGFR2-protein kinase C isoform epsilon signaling to increase the glycolytic metabolite, dihydroxyacetone.^19^ The Notch co-activator, forkhead box subfamily O1 transcription factor, couples vascular growth with metabolic activity, while Notch signaling triggers the phosphatidylinositol 3-kinase/AKT serine/threonine kinase pathway and regulates glycolysis-related genes during tumor angiogenesis.^63,64^

WSS is known to shape the formation of microvascular networks, allowing superfluous vessels to be pruned,^65^ and promoting angiogenesis at sites of high flow.^66^ WSS is implicated in reperfusion following injury,^38,39^ but the mechanism remains elusive. We expect reduction in blood flow in the immediate period following injury and in damaged vessels,^67^ and though vasomotor responses may be able to later augment flows to sites of injury,^68^ it is not clear how WSS signals can target growth to sites of injury. Here, we show that the geometry of the zebrafish trunk vasculature, in particular, a direct anastomotic connection between artery and vein-flow, is focused into a single Se vessel following injury. In spite of an overall decrease in flow, the robust peak WSS increase in this vessel turns it into a focus for reconnection of the network.

Arteriovenous anastomoses (“shunts”) can be observed across systems and organs,^69^ yet their existence is difficult to explain on the basis of efficient perfusion, since they offer red blood cells a high conductance route through tissues that circumvents the capillary bed. The anastomoses have been speculated to play a role in pressure and temperature regulation.^69^ In the zebrafish trunk, the anastomosis between DA and PCV is directly responsible for the repartitioning of flow following amputation, and thus for generating the WSS signals that initiate sprouting. Follow-up work should examine whether anastomoses in other systems provide the same function of shaping growth following injury, and whether the presence of anastomoses is correlated with robustness to damage.

## Methods

### The transgenic zebrafish tail amputation model for vascular injury and regeneration

Zebrafish embryos were harvested from natural mating at the UCLA Zebrafish Core Facility. The transgenic *Tg(fli1: gfp)* line, in which endothelial vasculature displayed GFP under the control of tissue specific *Fli1* promoter (ERGB) was used to assess vascular injury and regeneration. Zebrafish embryos at 1-2 cell stage of development were collected for micro-injections. Anti-sense MO against the ATG site of *p53* (0.5-1.0 mM, GeneTools LLC, OR) was utilized as standard control against cytotoxic damage from MO injection.^70^ In addition to the control *p53* MO, *epo* mRNA (10-20 pg/nL) or *Gata1a* MO (1mM, GeneTools LLC, OR) were micro-injected respectively to manipulate the level of hematopoiesis and subsequent viscosity-mediated WSS. *NICD* or DN-*Notch1b* mRNAs (10-20 pg/nL) were used to modulate global Notch activation. Anti-sense *ephrinb2a* & *EphB4* MOs (0.5-1.0 mM, GeneTools LLC, OR) and custom-designed *ephrinb2* mRNA (10-20 pg/nL) were micro-injected to manipulate global ephrinb2 expression.

Immediately after micro-injection, embryos were cultivated at 28.5 °C for 3 days in fresh standard E3 medium supplemented with 0.05% methylene blue (Sigma Aldrich, MO) and 0.003% phenylthiourea (PTU, Sigma Aldrich, MO) to suppress fungal outbreak and prevent melanogenesis. At 3 dpf, embryos were randomly selected for tail amputation as previously described.^71^ Following tail amputation, embryos were micro-injected with 6% hydroxyethyl hetastarch (Sigma Aldrich, MO), returned to fresh E3 medium, or E3 medium dosed with pharmacological inhibitors. These inhibitors included γ-secretase inhibitor (DAPT, 100 µM), isoproterenol (100 µM) and BDM (100 µM, Sigma Aldrich, MO). At 4 dpa, dual channel confocal imaging was performed to assess vessel regeneration as previously described.^9^ Z-scanned images were projected in the visualization plane where voxels displayed maximum intensity. **Table 1** provides the sequences of all MOs used in this study.

**Table 1.**
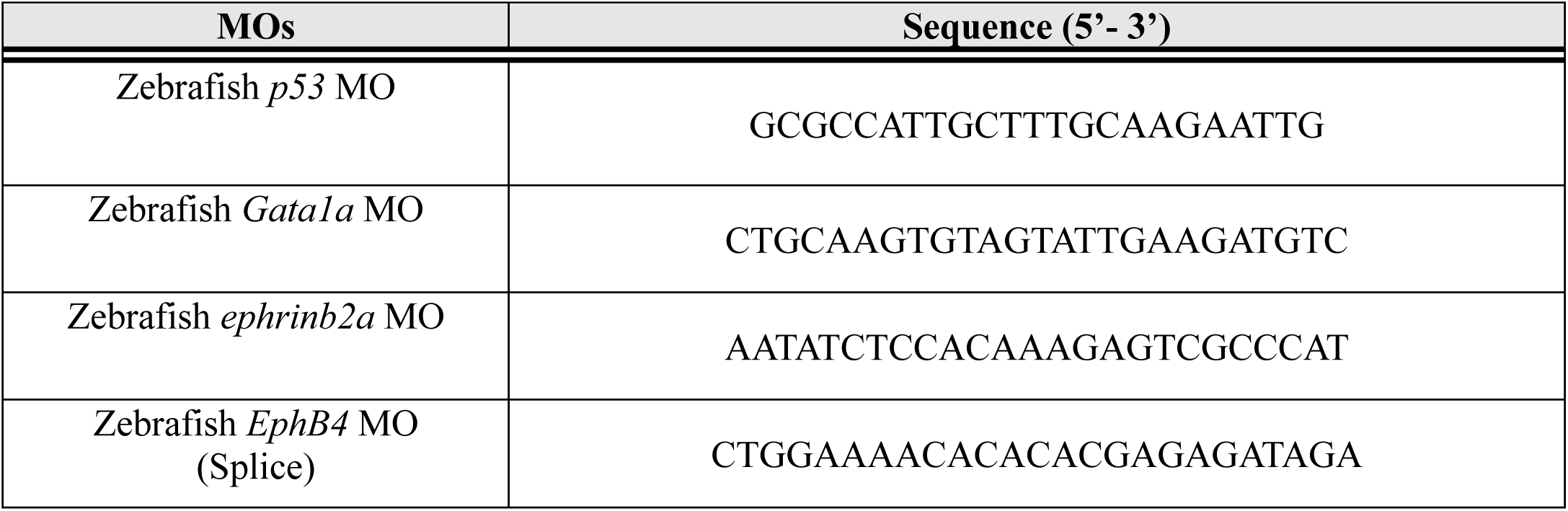
Sequencing Information of MOs.

### 6% hydroxyethyl hetastarch injection via common cardinal vein (CCV)

Double transgenic *Tg(fli1:gfp; gata1:ds-red)* embryos were immobilized with neutralized tricaine (Sigma Aldrich, MO) in 3% agarose to perform 6% hydroxyethyl hetastarch injection. The CCV and injection site were located anatomically. The injection was recapitulated under inverted immunofluorescence microscope (Olympus, IX70) by co-injecting fluorescein isothiocyanate conjugated dextran (FITC-dextran, Sigma Aldrich, MO). **Supplemental Figure 6A** shows distribution of FITC-dextran post 6% hydroxyethyl hetastarch injection. Average heart rate was assessed manually at 1 hour post injection. To quantify changes in volume and plasma viscosity, sequential images of aortic flow (*ds-red*^*+*^) were processed to generate binary images by using ImageJ (NIH, MD). Red and blue boxes in **Supplemental Figure 6C** depict regions of interest to measure variations of viscosity (%, total length of *ds-red*^*+*^ per unit length of DA, red boxes) and flow rate (numbers of *ds-red*^*+*^ per unit length of DA, blue boxes) following 6% hydroxyethyl hetastarch injection. By using cultured HAEC, endogenous expression of Notch-related genes under static condition was assessed following 4 and 12 hours of 6% hydroxyethyl hetastarch treatment.

### Assessment of spatiotemporal variations in endothelial *tp1* activity

The transgenic *Tg(tp1: gfp)* line was crossbred with the *Tg(flk1:mCherry)* line to visualize the activity of the Rbp-Jκ responsive element (Epstein Barr Virus terminal protein 1, *tp1*) in the vascular endothelial network. In response to genetic and pharmacologic manipulations of WSS or global Notch activity, spatiotemporal variations in endothelial *tp1* were sequentially imaged with dual channel confocal microscopy (Leica SP8, Germany) for 4 consecutive days post tail amputation. Acquired images were superimposed and analyzed by ImageJ.

### Whole mount zebrafish Immunofluorescence staining

Following tail amputation, *Tg(flk1: mCherry)* zebrafish embryos were fixed in 10% neutral buffered formalin solution (Sigma Aldrich, MO), dehydrated in methanol (Thermofisher, MA) and permeabilized in ice cold pure acetone (Sigma Aldrich, MO). Following permeabilization, zebrafish embryos were washed and blocked with 3% bovine serum albumin (Sigma Aldrich, MO) in 0.2% PBST (PBS + Triton-X, Sigma Aldrich, MO). Primary antibodies anti-collagen 4 (Ab6586, Abcam), anti-ephrinb2 (Ab150411, Abcam), and anti-phosphorylated histone H3 (Ser10) (06-570, Sigma Aldrich, MO) were used to detect corresponding protein expression and cell proliferation.^72^ Following overnight incubation, anti-rabbit (ab150077, Abcam) or anti-mouse IgG (ab150113, Abcam) conjugated to Alexa-488 was used to amplify primary-specific fluorescence.

### Quantitative real-time polymerase chain reaction (qRT-PCR) analyses

Total RNA was purified with Bio-Rad total RNA kit (Bio-Rad, CA) and reverse-transcribed to complementary DNA (cDNA) using a iScript cDNA synthesis kit (Bio-Rad, CA).^19^ Polymerase Chain Reaction was performed using qPCR master-mix (Applied Biological Materials Inc., Canada). The primer sequences are listed in **Table 2**. The expression of individual target mRNAs was normalized to human actin expression.

**Table 2.**
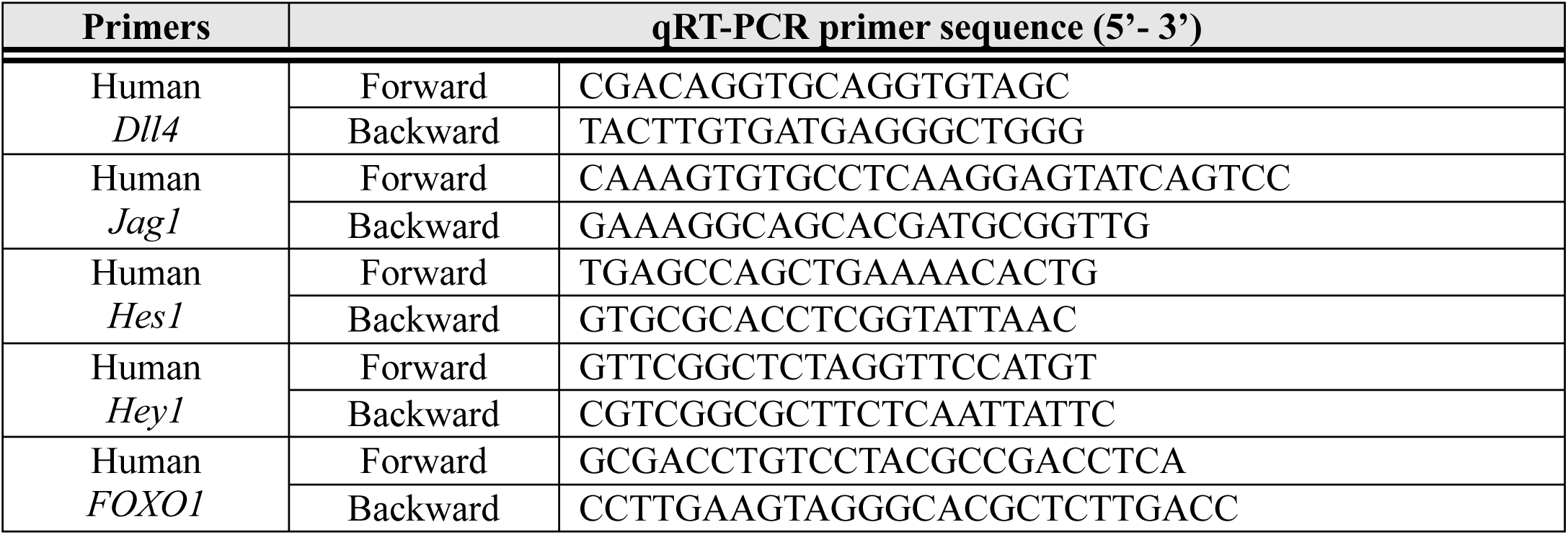
Sequencing Information of qRT-PCR primers.

### Batch processing of fluorescence images to examine endothelial-specific expression

To quantify co-localizations of fluorophores, multi-level thresholds based on Otsu’s method were used to segment single image slices in each fluorescent channel.^73^ The threshold level for each channel were manually selected to extract the most accurate binary masks of each slice. Pixels that represented vascular endothelium (*flk1*^*+*^) and the protein of interest were flagged with a value of 1, while the remaining region was regarded as background and flagged with a value of 0. Two channels of corresponding segmented images were merged together to generate the overlapping masks. The **Supplemental Figure 4A** depicts a schematic representation of the current method. To reduce intrinsic autofluorescence from the tissue, we pre-defined the effective area of mask from 20 to 500 pixels, that is, from 25.8 to 1290 μm^2^. The image post-processing, segmentation and rendering were processed by MATLAB (Mathworks, MA) and ImageJ.

### Imaging blood flow to assess the formation of vascular lumen during regeneration

An inverted immunofluorescence microscope (Olympus, IX70) and digital CCD camera (QIclick, Teledyne Qimaging, Canada) was used to sequentially image vascular injury and regeneration in the presence of blood flow. Images were superimposed by using ImageJ and Corel Imaging Software (ON, Canada).

### Measuring blood flow velocity and viscosity

To measure the blood flow velocity and viscosity in each Se vessel, we took image sequences at 20-40 frames per second, and for each sequence manually traced the center line of each Se vessel, without distinguishing between Se arteries and veins. We analyzed vessel flows using codes custom written in MATLAB. The code detects *ds-red*^*+*^ in each vessel by locating points with peak intensity. Peaks that are closer than the radius of the *ds-red*^*+*^ (∼3 μm) are coalesced into one cell at the centroid of the coalesced peaks. The number density of *ds-red*^*+*^ is calculated by dividing the number of *ds-red*^*+*^ by the length of the vessel. To obtain the velocity, *ds-red*^*+*^ detection is carried out in two consecutive frames, and for each *ds-red*^*+*^ detected in the first frame, the closest *ds-red*^*+*^ in the second frame is identified. If no *ds-red*^*+*^ is found within 30 μm, the velocity of the *ds-red*^*+*^ is not calculated. Since blood flow can only have one direction in a Se vessel, after *ds-red*^*+*^ velocities in all frames are calculated, the flow direction of each vessel is determined by majority rule, and *ds-red*^*+*^ velocities counter to the overall flow direction are removed.

### Endothelial WSS calculation

To calculate the WSS in each Se vessel, we separated regions adjacent to detected *ds-red*^*+*^ from the rest of the vessel. For the parts of the Se vessel that do not contain an *ds-red*^*+*^ the WSS is

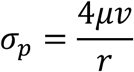

where *μ* is the viscosity, *v* is the median velocity of any detected *ds-red*^*+*^, and *r* is the radius of the vessel. For the part of the vessel adjacent to *ds-red*^*+*^, we use a model that increments the total resistance of the vessel by a constant, *α*_*c*_, for each *ds-red*^*+*^ contained in the vessel.^22^ The occlusive strength gives an increment on vessel resistance for each *ds-red*^*+*^ in the vessel. From force balance between pressure drop across the vessel and the shear force, and the shear force from the plasma part of the vessel given by the Poiseuille flow, the shear stress of the *ds-red*^*+*^ part of the vessel is

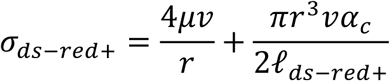

*𝓁*_*ds*−*red*+_ is the length of *ds-red*^*+*^. In this context, we used occlusive strengths *α*_*c*_ previously measured for 4 dpf wild type fish,^22^ which decreases from the mid-trunk to the tail. We use a uniform radius of 3 μm across all Se vessels, *𝓁*_*ds*−*red*+_ = 6*μm*, and μ=10^−3^ Pa•s.

### Quantification of vascular regeneration

Vessel segmentation and area quantification of the regenerated vascular loop were performed using Amira™ 3D imaging software (Thermofisher, MA). **Supplemental Figure 12** shows representative images. The entire vascular network in the posterior tail segment (purple) was segmented automatically, whereas a loop of regenerated vessel (pink) was assessed manually. The number of pixels was evaluated for statistical comparison.

### Preparation of mRNAs for *in vivo* rescue experiments

Rat *NICD*, zebrafish *epo* & DN-*Notch1b* cDNA were prepared as previously described.^8,71^ For transient ectopic overexpression of *ephrinb2* mRNA, zebrafish ephrinb2 cDNA was amplified from zebrafish cDNA with primers and cloned into pCS2^+^ at EcoRI/Xhol sites. Clones containing the insert were selected by PCR screening, and the clones were validated by sequencing. *In vitro* transcription was performed by using mMessage SP6 kit (Thermofisher, MA). For *in vivo* rescue experiments, transcribed mRNAs were purified by using a total RNA isolation kit (Bio-Rad, CA). The sequences of cloning primers are listed in **Table 3**.

**Table 3.**
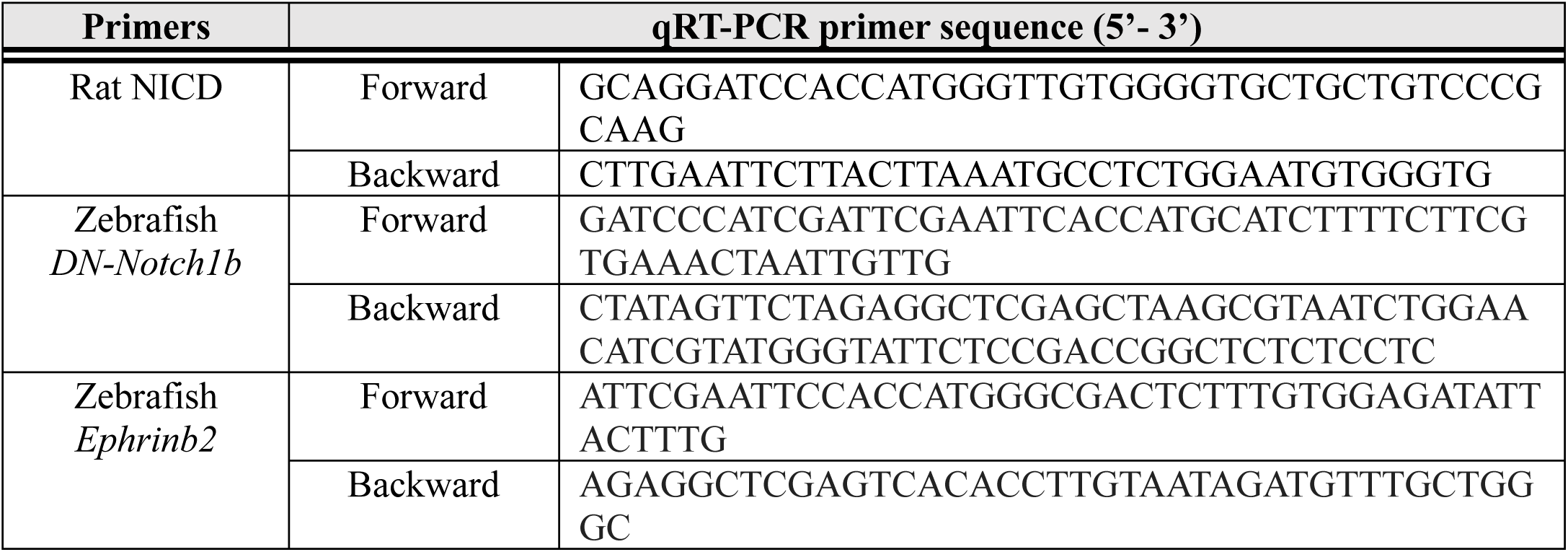
Cloning Primers.

### Small interference RNA (siRNA) transfection to cultured endothelial cells (EC)

Primary HAEC (Cell Applications, CA) were grown on bovine gelatin (Sigma Aldrich, MO)-coated-plates (Midsci, MO) at 37 °C and 5% CO2 and propagated for experiments between passages 4 and 10. EC growth medium (Cell Applications, CA) was supplemented with 5% fetal bovine serum (FBS, Life technologies, NY) and 1% penicillin-streptomycin (Life Technologies, NY) for optimal EC cultivation. At ∼50% confluency, FlexiTube™ siRNAs targeting scrambled negative control (Scr), Notch1, or ephrinb2/ EphB4 (Qiagen, Germany) were transfected following manufacturer’s instruction. Lipofectamine™ RNAiMAX (Thermofisher, MA) diluted with dulbecco’s modified eagle medium (DMEM)/10% FBS and Opti-MEM media with reduced serum (Thermofisher, MA) was used for siRNA transfection. Efficacy of transfections was verified by immunoblotting.

### EC migration and Matrigel tube formation assay

HAEC migration assay was performed as previously described.^9^ Areas between inner borders of HAEC at 4, 6, 12, 24 hours post scratch were evaluated using ImageJ. **Supplemental Figure 6F** shows a schematic representation of how HAEC migration was quantified. A tube formation assay was performed by seeding HAEC in 96-well plate coated with Matrigel with reduced growth factors (BD Biosciences, CA) in the presence of human VEGF (10-20 ng/mL, Sigma Aldrich, MO). Following 4 hours of incubation at 37 °C, tube formation was evaluated under an Olympus IX 70 phase-contrast microscope. The number of branching points and tube length were quantified manually using ImageJ.

### PSS exposure

Confluent monolayers of HAEC grown on 6-well plate were exposed to unidirectional PSS (6, 12 and 24 hours) using a modified flow device.^10^ Neutralized MCDB-131 medium (Sigma Aldrich, MO) containing 7.5% sodium bicarbonate solution, 10% FBS and 4% dextran from *Leuconostoc spp* (Sigma Aldrich, MO) was used for PSS exposure. Following exposure to PSS, the center of each monolayer was removed by using a cell scraper to collect only flow-aligned cells from the periphery of the well. To visualize changes in endogenous protein expressions and proximity ligations, we utilized our in-house dynamic flow system (*∂*τ/*∂*t = 29.3 dyne·cm^-2^·s^-1^, with time-averaged shear stress = 50 dyne·cm^-1^ at 1 Hz).^74^

### Immunoprecipitation, proximity ligation assays, and immunoblot analysis

Following PSS exposure, flow-aligned cells were lysed with M-PER mammalian protein extraction reagent (Thermofisher, MA) supplemented with 1% protease and phosphatase inhibitor cocktail (Thermofisher, MA) at 4°C. Detergent compatible protein assay (Bio-rad, CA) and lithium dodecyl sulfate polyacrylamide gel electrophoresis (Invitrogen, CA) were performed as previously described.^75^ Primary antibodies anti-Notch1 (MA5-32080, Thermofisher, MA), anti-ephrinb2 (Ab150411, Abcam), and anti-EphB4 (H-200, Santa Cruz Biotechnology Inc., TX) were used. Equal loading was verified by using anti-β-tubulin (AA2, Santa Cruz Biotechnology Inc., TX). Anti-rabbit (7074S, Cell Signaling Technology) or anti-mouse IgG (7076S, Cell Signaling Technology) conjugated to horseradish peroxidase was used for the secondary incubation.

Immunoprecipitation against ephrinb2/EphB4 was performed using a Pierce™ Crosslink magnetic IP/Co-IP kit (Thermofisher, MA). Disuccinmidiyl suberate was used to crosslink anti-ephrinb2, neutralized for immunoblot analyses. Densitometry was performed as previous described.^19^ Proximity ligations between ephrinb2/EphB4 in flow-aligned cells were conducted by using Duolink^®^ In situ Red Starter Kit Mouse/Rabbit (Sigma Aldrich, MO). Numbers of individual ligations in raw and post-processed images were compared to test the validity of the quantification. To evaluate polarization kinetics and distributions of the ligations, platelet endothelial adhesion molecule 1 (H-3, Santa Cruz Biotechnology Inc., TX) and 4′,6-diamidino-2-phenylindole (SC-3598, Santa Cruz Biotechnology Inc., TX) were fluorescently labeled.

### Statistics

Data were expressed as mean ± standard deviation and compared among separate experiments. Unpaired two-tail *t* test and 2-proportion z-test were used for statistical comparisons between 2 experimental conditions. *P* values < 0.05 were considered significant. Comparisons of multiple values were made by one-way analysis of variance (ANOVA) and statistical significance for pairwise comparison was determined by using the Tukey test.

### Study approval

Zebrafish experiments were performed in compliance with the Institutional Animal Care and Use Committees (IACUC) at the University of California, Los Angeles (UCLA), under animal welfare assurance number A3196-01.

## Supporting information

Supplemental Video 1 unamputated

Supplemental Video 1 amputated

Supplemental Video 2 0 dpa

Supplemental Video 2 2 dpa

Supplemental Video 2 4 dpa

## Author contributions

KIB and CCC performed zebrafish studies including micro-injections and confocal imaging. KIB, SSC, SC, MR, TKH wrote the manuscript. KIB, SSC, YW performed blood flow imaging and mathematical analyses of peak WSS. MR and JC performed CFD analyses. KIB, CCC and YD performed post image processing and batch analyses. KIB, JJM and JWA performed in vitro cell culture, flow exposure and molecular biology. KIB designed experiments. XX, HC, RO, SC, MR, and TKH supervised, revised, and supported the study.

## Acknowledgements

The authors like to express their gratitude to Dr. Weinmaster for providing the NICD plasmid, Drs. David Traver and Deborah Yellon at UCSD, Nathan Lawson at the University of Massachusetts Medical School, and Dr. Schulte-Merker at University of Münster for generously providing *Tg(tp1:GFP)* and *Tg(flt1:tdtomatoe; flt4;yfp)* zebrafish line. This study was supported by National Institutes of Health R01HL083015 (TKH), R01HL111437 (TKH), R01HL129727 (TKH), R01HL118650 (TKH), I01 BX004356 (TKH), 5R01GM126556 (MR), K99HL148493 (YD), and AHA 18CDA34110338 (YD).

## Author Disclosure Statement

All authors declare no competing financial interests.

## Supplemental Figure Legends

**Supplemental figure 1. Average diameter of segmental vessels in the amputated site.** Amputated vascular networks in *Tg(fli1: gfp)* zebrafish embryos were imaged to assess vessel diameter in SeAs and SeVs for flow adaptation. Average diameters of the caudal SeAs adjacent to the amputated site modestly increased from 1 dpa and remained dilated during regeneration.

**Supplemental figure 2. Computational fluid dynamics (CFD) to validate hemodynamic WSS modulation** CFD analyses were performed to validate viscosity- and contractility-mediated hemodynamic WSS. While increase in viscosity (*epo* mRNA, 10-20 pg/nL) or myocardial contractility (isoproterenol, 100 μM) increased averaged hemodynamic WSS at 0 dpa, reduction of viscosity (*Gata1a* MO, 1 mM) or myocardial contractility (BDM, 100 μM) reduced WSS in the distal SeA adjacent to the amputated site (white arrow). SeA: Arterial segmental vessel, SeV: Venous segmental vessel.

**Supplemental figure 3. Maximum intensity projections of wholemount *pHH3***^***+***^ **in the amputated site** Cell mitosis marker (*pHH3*^*+*^) was fluorescently stained in the *Tg(flk1: mcherry)* line to quantify Notch-dependent proliferation. **(A)** Time-lapse image of the total *pHH3*^*+*^ cells with and without DAPT treatment. Scale bar: 20 μm. **(B)** Representative images of the total *pHH3*^*+*^ cells in response to differential hemodynamic WSS. Scale bar: 20 μm.

**Supplemental figure 4.** Schematic representations of batch processing and multi-level threshold analyses to visualize 3D colocalizations of two fluorophores.

**Supplemental figure 5. Endothelial Notch signaling regulates endothelial cell migration and tube formation *in vitro* (A-B)** Representative images of Matrigel tube formation and HAEC migration with or without DAPT treatment or *siNotch1* transfection (Black arrowheads). **(C)** The density quantification of Notch1 protein expressions following *siScr* or *siNotch1* transfection. TCL: total cell lysates. **(D-E)** DAPT treatment or *siNotch1* transfection reduced both tube length and the number of branch points as compared to DMSO-or *siScr-*transfected controls (^*\*\**^ *p < 0.005* vs. DMSO, vs. *siScr*, n=3). **(F)** DAPT treatment or *siNotch1* transfection significantly attenuated area of recovery (A.O.R) at 12 hours post scratch. (^*\**^ *p < 0.005*, ^*\*\**^ *p < 0.005* vs. DMSO, vs. *siScr*, n=3) **(G)** 6 hours of in vitro PSS exposure upregulated Notch-related gene expressions including Notch ligands Dll4 and Jag1, and the targets Hes1, Hey1 and Hey2. DAPT treatment mitigated PSS-increased mRNA expressions, whereas *siNotch1* transfection specifically inhibited Dll4-Hes1 axis (^*\**^ *p < 0.05*, ^*\*\**^ *p < 0.005*, ^*\*\*\**^ *p < 0.0005* vs. static, normalized with human actin, n=3).

**Supplemental figure 6. Analyses of 6% hydroxyethyl hetastarch injection via common cardial vein (CCV) (A)** The injection of 6% hydroxyethyl hetastarch to modulate plasma viscosity was recapitulated by co-injecting with FITC-dextran. Images of injected embryos were taken at 1 hour post injection (hpi). DLAV: Dorsal longitudinal anastomotic vessel. CCV: Common cardinal vein. Scale bar: 50 μm. **(B)** Average heart rate (bpm) remained unchanged at 1 hpi (n=5). **(C)** Assessment of changes in plasma viscosity and flow rate following 6% hydroxyethyl hetastarch injection. For the measurements, see Materials & Method. **(D-E)** At 1 hpi, viscosity in the DA was reduced by 5.5%, whereas flow rate of *ds-red*^*+*^ remained the unchanged (n=6 for viscosity, n=4 for flow rate measurement). **(F)** Under the static condition, Notch-related gene expressions in HAEC remained unchanged following 4 and 12 hours of 6% hydroxyethyl hetastarch treatment (^*\*\**^ *p < 0.005*, ^*\*\*\**^ *p < 0.0005* vs. H2O, normalized to human actin, n=3).

**Supplemental figure 7. Notch signaling pathway regulates WSS-mediated loop formation**

**(A)** Representative images of endothelial *tp1* activity and vessel regeneration following transient modulations of global Notch expressions (DAPT, DN-*Notch1b* and *NICD* mRNA) and WSS (*epo* mRNA, isoproterenol, *Gata1a* MO and BDM). In the presence of DAPT or DN-*Notch1b* mRNA, augmented WSS failed to increase endothelial *tp1* activity in the amputated site (white arrowheads) to promote loop formation at 4 dpa (white arrows). *NICD* mRNA injection up-regulated endothelial *tp1* activity and restored *Gata1a* MO-impaired regeneration. Following exposure to BDM, *NICD* mRNA increased endothelial *tp1* activity, but failed to restore regeneration. (^*\**^ *p < 0.05*, ^*\*\**^ *p < 0.005*, vs. *NICD* mRNA+*Gata1a* MO, n=20).

**Supplemental figure 8. Time-lapse imaging of arterial and venous vessels with *Tg(Flt1:tdtomatoe; Flt4: yfp)* zebrafish** Time-lapse images of arterial (*flt1*^*+*^, red) and venous (*flt4*^+^, green) vessels with indicated treatments.

**Supplemental figure 9. Notch signaling pathway regulates WSS-responsive arterial network** In the presence of DAPT or DN-*Notch1b* mRNA, increase in viscosity- and contractility-mediated WSS (*epo* mRNA, isoproterenol, 6% hydroxyethyl hetastarch) resulted in the absence of *flt1*^*+*^network (white arrowheads) for loop formation (white arrow) at 4 dpa. In addition, *flt4*^*+*^ DLAV and PCV (* asterisk) was partially attenuated at 4 dpa. Conversely, *NICD* mRNA reversed the effect of *Gata1a* MO (n=3 per each time point).

**Supplemental figure 10. Endothelial ephrinb2/Ephb4 pathway systematically regulates vascular loop formation (A)** Injection of EphB4 MO (0.5 mM) disrupted the CVP morphology (*asterisk) and impaired DLAV-PCV loop formation (white arrowheads). In addition, knockdown of both ephrinb2 and EphB4 (0.25 mM) sustained EphB4-knockdown phenotypic patterns and arrested loop formation (n=20 per each group) EphB4 or double knockdown inhibited both *flt1*^*+*^ in the proximal SeA and *flt4*^*+*^ DLAV and PCV during loop formation. (n=20 per each group) **(B)** Schematic representations of ephrinb2/EphB4-mediated arterial and venous regeneration. Black arrowheads depict regenerated *flt1*^*+*^/*flt4*^+^ following EphB4 or ephrinb2/EphB4 knockdown. **(C-D)** Representative images of Matrigel tube formation and HAEC migration with and without *siEphB4* transfection and/or co-transfection with *siephrinb2*. **(E)** The density quantification of EphB4 expression following *siEphB4* transfection. TCL: total cell lysate (**F-G**) While *siEphB4* transfection reduced tube length and the number of branch points, transfection of both *siephrinb2* and *siEphB4* aggravated branch point formation as compared to *siScr*-transfected HAEC. (^*\**^ *p < 0.05*, ^*\*\**^ *p < 0.005*, ^*\*\*\**^ *p < 0.0005* vs. *siScr*, n=3) **(H)** *siEphB4* transfection reduced, whereas transfection of both *siephrinb2* and *siEphB4* further reduced area of recovery (A.O.R) by at 24 hours post scratch (^*\*\**^ *p < 0.005*, ^*\*\*\**^ *p < 0.0005* vs. vs. *siScr*, n=3).

**Supplemental figure 11. Pulsatile shear stress (PSS) increases total amount of ephrinb2/EphB4 interaction** Human Aortic Endothelial Cells (HAEC) monolayers were subjected to unidirectional PSS (23±7 dyne·cm^-2^ at 1Hz) for 6, 12, 24 hours. **(A)** Exposure to PSS increased both ephrinb2 and EphB4 protein expression in a time-dependent manner. While DAPT treatment and *siNotch1* transfection attenuated PSS-increased ephrinb2 expressions, EphB4 expression remained statistically unchanged. The density quantification of the blots was normalized to β-tubulin. (n=3) **(B)** Transcript of klf2 mRNA expressions were assessed as an internal control. klf2 mRNA increased by ∼ 4.0-fold following PSS exposure. (^*\**^ *p < 0.05*, vs. static, normalized to human actin, n=3) **(C)** *siephrinb2* transfection did not affect PSS-increased EphB4. **(D)** Compared to static controls, 6 hours of PSS exposure modulated ephrinb2-mediated EphB4 pull down in Notch-dependent manner (untreated: 59%; *siScr*: 65%, *siNotch1*: 18% reduction, DMSO: 21% and DAPT: 16%, respectively, ^*\**^ *p < 0.05*, vs. static, n=3). **(E-F)** PSS further increased the total amount of ephrinb2/EphB4 complex without affecting polarization kinetics between endogenous ephrinb2 and EphB4. (^*\**^ *p < 0.05* vs. static, n=3) The average number of individual ephrinb2/EphB4 ligation and average size of the total ligations per cell were quantified for statistical comparisons. Compared to the static control, PSS exposure increased average size of an individual ligation by 2.3-fold, while *siNotch1* transfection significantly reduced both the number and the size of ligations and attenuated the effect of PSS (^*\**^ *p < 0.05* vs. static, n=3).

**Supplemental figure 12. Quantification of vascular loop formation**

Area quantifications of the vessel regeneration were performed as described in Method. Entire network of endothelial vasculature on the posterior tail segment (purple) was designated automatically, whereas regenerated vascular loop was derived manually (pink).

## Supplemental Videos

**Supplemental Videos 1. Tail amputation increased hemodynamic flow in the distal segmental arteries (SeA)**

**Supplemental Videos 2. Blood flow adaptation in *flt1***^***+***^ **network following tail amputation.**

## Graphic Abstract

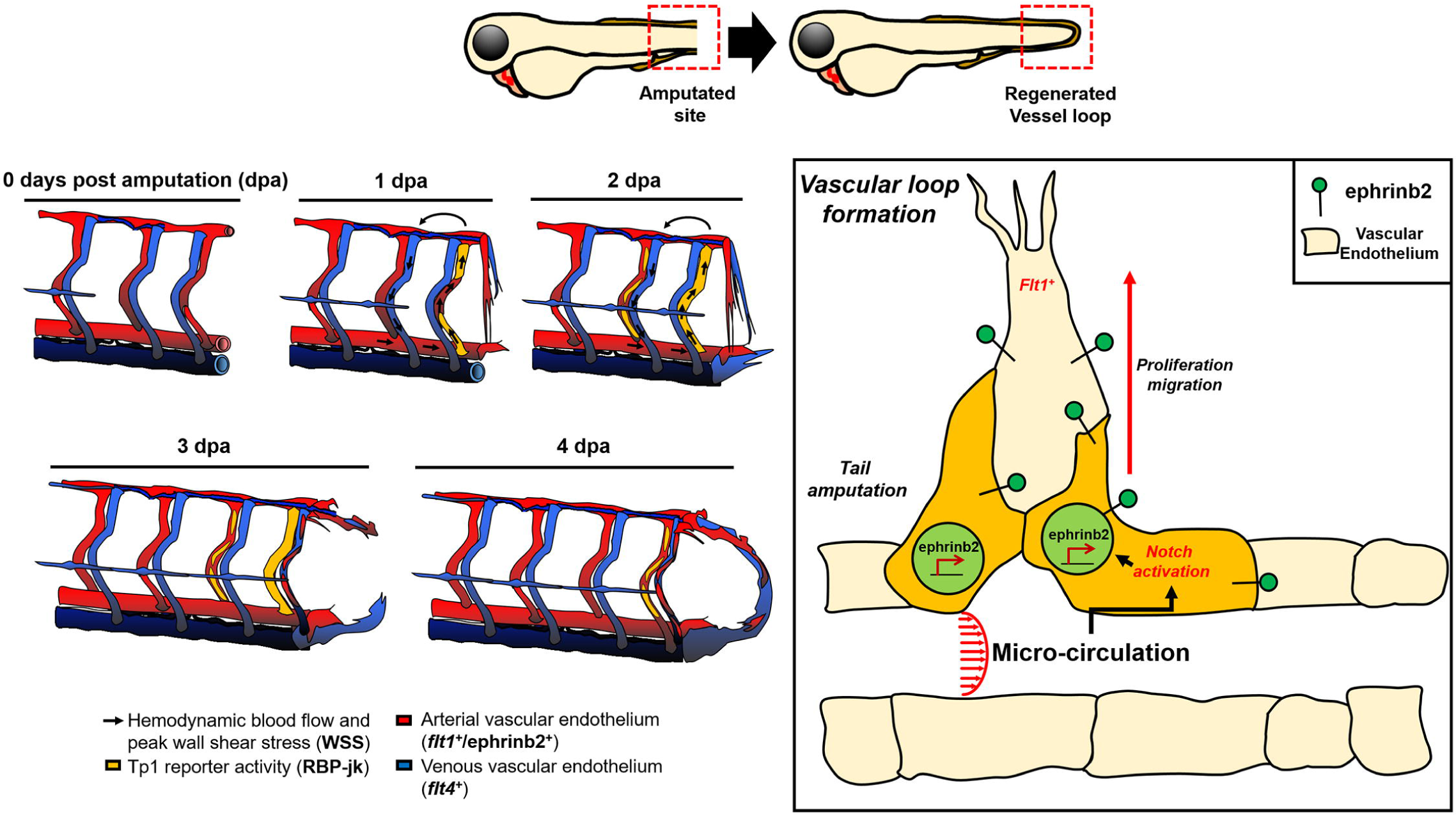

## Notes

### Competing Interest Statement

The authors have declared no competing interest.

## References

1. Dahl A, Cai N, Ko A, et al. Reverse GWAS: Using genetics to identify and model phenotypic subtypes. PLoS genetics. 2019;15(4):e1008009.

2. Ahlqvist E, Storm P, Käräjämäki A, et al. Novel subgroups of adult-onset diabetes and their association with outcomes: a data-driven cluster analysis of six variables. The lancet Diabetes & endocrinology. 2018;6(5):361–369.

3. Topper JN, Gimbrone MA, Jr. Blood flow and vascular gene expression: fluid shear stress as a modulator of endothelial phenotype. Molecular medicine today. Jan 1999;5(1):40–46.

4. Tanaka H, Shimizu S, Ohmori F, et al. Increases in blood flow and shear stress to nonworking limbs during incremental exercise. Medicine and science in sports and exercise. Jan 2006;38(1):81–85.

5. Loerakker S, Stassen O, Ter Huurne FM, Boareto M, Bouten CVC, Sahlgren CM. Mechanosensitivity of Jagged-Notch signaling can induce a switch-type behavior in vascular homeostasis. Proceedings of the National Academy of Sciences of the United States of America. Apr 17 2018;115(16):E3682–E3691.

6. Masumura T, Yamamoto K, Shimizu N, Obi S, Ando J. Shear stress increases expression of the arterial endothelial marker ephrinB2 in murine ES cells via the VEGF-Notch signaling pathways. Arteriosclerosis, thrombosis, and vascular biology. 2009;29(12):2125–2131.

7. Bray SJ. Notch signalling in context. Nature reviews. Molecular cell biology. Nov 2016;17(11):722–735.

8. Lee J, Fei P, Packard RR, et al. 4-Dimensional light-sheet microscopy to elucidate shear stress modulation of cardiac trabeculation. J Clin Invest. May 2 2016;126(5):1679–1690.

9. Lee J, Vedula V, Baek KI, et al. Spatial and temporal variations in hemodynamic forces initiate cardiac trabeculation. JCI insight. 2018;3(13).

10. Mack JJ, Mosqueiro TS, Archer BJ, et al. NOTCH1 is a mechanosensor in adult arteries. Nature communications. Nov 20 2017;8(1):1620.

11. Adams RH, Wilkinson GA, Weiss C, et al. Roles of ephrinB ligands and EphB receptors in cardiovascular development: demarcation of arterial/venous domains, vascular morphogenesis, and sprouting angiogenesis. Genes & development. 1999;13(3):295–306.

12. Gerety SS, Anderson DJ. Cardiovascular ephrinB2 function is essential for embryonic angiogenesis. Development. 2002;129(6):1397–1410.

13. Wilkinson DG. Eph receptors and ephrins: regulators of guidance and assembly. International review of cytology. Vol 196: Elsevier; 2000:177–244.

14. Frisén J, Holmberg J, Barbacid M. Ephrins and their Eph receptors: multitalented directors of embryonic development. The EMBO journal. 1999;18(19):5159–5165.

15. Kullander K, Klein R. Mechanisms and functions of Eph and ephrin signalling. Nature reviews Molecular cell biology. 2002;3(7):475–486.

16. Lawson ND, Scheer N, Pham VN, et al. Notch signaling is required for arterial-venous differentiation during embryonic vascular development. Development. 2001;128(19):3675–3683.

17. Swift MR, Weinstein BM. Arterial–venous specification during development. Circulation research. 2009;104(5):576–588.

18. Füller T, Korff T, Kilian A, Dandekar G, Augustin HG. Forward EphB4 signaling in endothelial cells controls cellular repulsion and segregation from ephrinB2 positive cells. Journal of cell science. 2003;116(12):2461–2470.

19. Baek KI, Li R, Jen N, et al. Flow-Responsive Vascular Endothelial Growth Factor Receptor-Protein Kinase C Isoform Epsilon Signaling Mediates Glycolytic Metabolites for Vascular Repair. Antioxid Redox Signal. Jan 1 2018;28(1):31–43.

20. Tzahor E, Poss KD. Cardiac regeneration strategies: staying young at heart. Science. 2017;356(6342):1035–1039.

21. Zhao L, Ben-Yair R, Burns CE, Burns CG. Endocardial Notch signaling promotes cardiomyocyte proliferation in the regenerating zebrafish heart through Wnt pathway antagonism. Cell reports. 2019;26(3):546-554. e545.

22. Chang S-S, Tu S, Baek KI, et al. Optimal occlusion uniformly partitions red blood cells fluxes within a microvascular network. PLoS computational biology. 2017;13(12):e1005892.

23. Sugden WW, Meissner R, Aegerter-Wilmsen T, et al. Endoglin controls blood vessel diameter through endothelial cell shape changes in response to haemodynamic cues. Nature cell biology. 2017;19(6):653.

24. Chouinard-Pelletier G, Jahnsen ED, Jones EA. Increased shear stress inhibits angiogenesis in veins and not arteries during vascular development. Angiogenesis. 2013;16(1):71–83.

25. Kametani Y, Chi NC, Stainier DY, Takada S. Notch signaling regulates venous arterialization during zebrafish fin regeneration. Genes to Cells. 2015;20(5):427–438.

26. Krebs LT, Xue Y, Norton CR, et al. Notch signaling is essential for vascular morphogenesis in mice. Genes Dev. Jun 1 2000;14(11):1343–1352.

27. Baonza A, Garcia-Bellido A. Notch signaling directly controls cell proliferation in the Drosophila wing disc. Proceedings of the National Academy of Sciences of the United States of America. Mar 14 2000;97(6):2609–2614.

28. Jensen J, Heller RS, Funder-Nielsen T, et al. Independent development of pancreatic alpha- and beta-cells from neurogenin3-expressing precursors: a role for the notch pathway in repression of premature differentiation. Diabetes. Feb 2000;49(2):163–176.

29. Fre S, Huyghe M, Mourikis P, Robine S, Louvard D, Artavanis-Tsakonas S. Notch signals control the fate of immature progenitor cells in the intestine. Nature. Jun 16 2005;435(7044):964–968.

30. Stanger BZ, Datar R, Murtaugh LC, Melton DA. Direct regulation of intestinal fate by Notch. Proceedings of the National Academy of Sciences of the United States of America. Aug 30 2005;102(35):12443–12448.

31. Pellegrinet L, Rodilla V, Liu Z, et al. Dll1- and dll4-mediated notch signaling are required for homeostasis of intestinal stem cells. Gastroenterology. Apr 2011;140(4):1230-1240 e1231-1237.

32. Artavanis-Tsakonas S, Rand MD, Lake RJ. Notch signaling: cell fate control and signal integration in development. Science. Apr 30 1999;284(5415):770–776.

33. Lobov IB, Renard RA, Papadopoulos N, et al. Delta-like ligand 4 (Dll4) is induced by VEGF as a negative regulator of angiogenic sprouting. Proceedings of the National Academy of Sciences of the United States of America. Feb 27 2007;104(9):3219–3224.

34. Kim YH, Hu H, Guevara-Gallardo S, Lam MT, Fong SY, Wang RA. Artery and vein size is balanced by Notch and ephrin B2/EphB4 during angiogenesis. Development. Nov 2008;135(22):3755–3764.

35. Corpechot C, Barbu V, Wendum D, et al. Hypoxia-induced VEGF and collagen I expressions are associated with angiogenesis and fibrogenesis in experimental cirrhosis. Hepatology. 2002;35(5):1010–1021.

36. Germain S, Monnot C, Muller L, Eichmann A. Hypoxia-driven angiogenesis: role of tip cells and extracellular matrix scaffolding. Current opinion in hematology. 2010;17(3):245–251.

37. Kochhan E, Lenard A, Ellertsdottir E, et al. Blood flow changes coincide with cellular rearrangements during blood vessel pruning in zebrafish embryos. PloS one. 2013;8(10).

38. Kaunas R, Kang H, Bayless KJ. Synergistic regulation of angiogenic sprouting by biochemical factors and wall shear stress. Cellular and molecular bioengineering. 2011;4(4):547–559.

39. Ichioka S, Shibata M, Kosaki K, Sato Y, Harii K, Kamiya A. Effects of shear stress on wound-healing angiogenesis in the rabbit ear chamber. Journal of Surgical Research. 1997;72(1):29–35.

40. Isogai S, Horiguchi M, Weinstein BM. The vascular anatomy of the developing zebrafish: an atlas of embryonic and early larval development. Developmental biology. 2001;230(2):278–301.

41. Fong G-H. Mechanisms of adaptive angiogenesis to tissue hypoxia. Angiogenesis. 2008;11(2):121–140.

42. Bochenek ML, Dickinson S, Astin JW, Adams RH, Nobes CD. Ephrin-B2 regulates endothelial cell morphology and motility independently of Eph-receptor binding. Journal of cell science. 2010;123(8):1235–1246.

43. Zheng LC, Wang XQ, Lu K, et al. Ephrin-B2/Fc promotes proliferation and migration, and suppresses apoptosis in human umbilical vein endothelial cells. Oncotarget. Jun 20 2017;8(25):41348–41363.

44. Månsson-Broberg A, Siddiqui AJ, Genander M, et al. Modulation of ephrinB2 leads to increased angiogenesis in ischemic myocardium and endothelial cell proliferation. Biochemical and biophysical research communications. 2008;373(3):355–359.

45. Wang Y, Nakayama M, Pitulescu ME, et al. Ephrin-B2 controls VEGF-induced angiogenesis and lymphangiogenesis. Nature. 2010;465(7297):483.

46. Hayashi S-i, Asahara T, Masuda H, Isner JM, Losordo DW. Functional ephrin-B2 expression for promotive interaction between arterial and venous vessels in postnatal neovascularization. Circulation. 2005;111(17):2210–2218.

47. Korff T, Braun J, Pfaff D, Augustin HG, Hecker M. Role of ephrinB2 expression in endothelial cells during arteriogenesis: impact on smooth muscle cell migration and monocyte recruitment. Blood, The Journal of the American Society of Hematology. 2008;112(1):73–81.

48. Bai J, Wang YJ, Liu L, Zhao YL. Ephrin B2 and EphB4 selectively mark arterial and venous vessels in cerebral arteriovenous malformation. The Journal of international medical research. Apr 2014;42(2):405–415.

49. Arora S, Lam AJY, Cheung C, Yim EKF, Toh YC. Determination of critical shear stress for maturation of human pluripotent stem cell-derived endothelial cells towards an arterial subtype. Biotechnology and bioengineering. May 2019;116(5):1164–1175.

50. Sivarapatna A, Ghaedi M, Le AV, Mendez JJ, Qyang Y, Niklason LE. Arterial specification of endothelial cells derived from human induced pluripotent stem cells in a biomimetic flow bioreactor. Biomaterials. 2015;53:621–633.

51. Augustin HG, Reiss Y. EphB receptors and ephrinB ligands: regulators of vascular assembly and homeostasis. Cell and tissue research. 2003;314(1):25–31.

52. Zhang G, Zhou J, Fan Q, et al. Arterial–venous endothelial cell fate is related to vascular endothelial growth factor and Notch status during human bone mesenchymal stem cell differentiation. FEBS letters. 2008;582(19):2957–2964.

53. Groppa E, Brkic S, Uccelli A, et al. EphrinB2/EphB4 signaling regulates non-sprouting angiogenesis by VEGF. EMBO reports. 2018;19(5).

54. Yang D, Jin C, Ma H, et al. EphrinB2/EphB4 pathway in postnatal angiogenesis: a potential therapeutic target for ischemic cardiovascular disease. Angiogenesis. 2016;19(3):297–309.

55. Fuller T, Korff T, Kilian A, Dandekar G, Augustin HG. Forward EphB4 signaling in endothelial cells controls cellular repulsion and segregation from ephrinB2 positive cells. J Cell Sci. Jun 15 2003;116(Pt 12):2461–2470.

56. Xu C, Hasan SS, Schmidt I, et al. Arteries are formed by vein-derived endothelial tip cells. Nature communications. 2014;5(1):1–11.

57. Chistiakov DA, Orekhov AN, Bobryshev YV. Effects of shear stress on endothelial cells: go with the flow. Acta physiologica. 2017;219(2):382–408.

58. Chistiakov DA, Orekhov AN, Bobryshev YV. The role of miR-126 in embryonic angiogenesis, adult vascular homeostasis, and vascular repair and its alterations in atherosclerotic disease. Journal of molecular and cellular cardiology. 2016;97:47–55.

59. Lou Y-L, Guo F, Liu F, et al. miR-210 activates notch signaling pathway in angiogenesis induced by cerebral ischemia. Molecular and cellular biochemistry. 2012;370(1-2):45–51.

60. Yang M, Li C-J, Sun X, et al. MiR-497∼ 195 cluster regulates angiogenesis during coupling with osteogenesis by maintaining endothelial Notch and HIF-1α activity. Nature communications. 2017;8(1):1–11.

61. Marcet B, Chevalier B, Luxardi G, et al. Control of vertebrate multiciliogenesis by miR-449 through direct repression of the Delta/Notch pathway. Nat Cell Biol. Jun 2011;13(6):693–699.

62. Wang W, Feng L, Zhang H, et al. Preeclampsia up-regulates angiogenesis-associated microRNA (ie., miR-17,-20a, and-20b) that target ephrin-B2 and EPHB4 in human placenta. The Journal of Clinical Endocrinology & Metabolism. 2012;97(6):E1051–E1059.

63. Landor SK, Mutvei AP, Mamaeva V, et al. Hypo- and hyperactivated Notch signaling induce a glycolytic switch through distinct mechanisms. Proceedings of the National Academy of Sciences of the United States of America. Nov 15 2011;108(46):18814–18819.

64. Slaninova V, Krafcikova M, Perez-Gomez R, et al. Notch stimulates growth by direct regulation of genes involved in the control of glycolysis and the tricarboxylic acid cycle. Open biology. Feb 2016;6(2):150155.

65. Chen Q, Jiang L, Li C, et al. Haemodynamics-driven developmental pruning of brain vasculature in zebrafish. PLoS biology. 2012;10(8).

66. Young S, Egginton S. AAllometry of skeletal muscle fine structure allows maintenance of aerobic capacity during ontogenetic growth. Journal of Experimental Biology. 2009;212(21):3564–3575.

67. Yamakami I, McIntosh TK. AAlterations in regional cerebral blood flow following brain injury in the rat. Journal of Cerebral Blood Flow & Metabolism. 1991;11(4):655–660.

68. Aulick LH, Baze WB, McLeod Jr CG, Wilmore DW. AControl of blood flow in a large surface wound. Annals of surgery. 1980;191(2):249.

69. Sherman JL. ANormal arteriovenous anastomoses. Medicine. 1963;42(4):247–268.

70. Robu ME, Larson JD, Nasevicius A, et al. p53 activation by knockdown technologies. PLoS Genet. May 25 2007;3(5):e78.

71. Baek KI, Packard RRS, Hsu JJ, et al. Ultrafine Particle Exposure Reveals the Importance of FOXO1/Notch Activation Complex for Vascular Regeneration. Antioxidants & redox signaling. 2018;28(13):1209–1223.

72. Elmaci I, Altinoz MA, Sari R, Bolukbasi FH. APhosphorylated Histone H3 (PHH3) as a Novel Cell Proliferation Marker and Prognosticator for Meningeal Tumors: A Short Review. Applied immunohistochemistry & molecular morphology : AIMM. Oct 2018;26(9):627–631.

73. Sezgin M, Sankur B. ASurvey over image thresholding techniques and quantitative performance evaluation. Journal of Electronic Imaging. 2004;13(1):146-165, 120.

74. Li R, Jen N, Wu L, et al. Disturbed Flow Induces Autophagy, but Impairs Autophagic Flux to Perturb Mitochondrial Homeostasis. Antioxid Redox Signal. Nov 20 2015;23(15):1207–1219.

75. Li R, Beebe T, Jen N, et al. Shear stress-activated Wnt-angiopoietin-2 signaling recapitulates vascular repair in zebrafish embryos. Arterioscler Thromb Vasc Biol. Oct 2014;34(10):2268–2275.

